# Whole Human-Brain Mapping of Single Cortical Neurons for Profiling Morphological Diversity and Stereotypy

**DOI:** 10.1101/2022.10.29.514375

**Authors:** Xiaofeng Han, Shuxia Guo, Nan Ji, Tian Li, Jian Liu, Xiangqiao Ye, Yi Wang, Zhixi Yun, Feng Xiong, Jing Rong, Di Liu, Hui Ma, Yujin Wang, Yue Huang, Peng Zhang, Wenhao Wu, Liya Ding, Michael Hawrylycz, Ed Lein, Giorgio A. Ascoli, Wei Xie, Lijuan Liu, Liwei Zhang, Hanchuan Peng

## Abstract

Quantification of individual cells’ morphology and their distribution at the whole brain scale is essential to understand the structure and diversity of cell types. Despite recent technological advances, especially single cell labeling and whole brain imaging, for many prevailing animal models, it is exceedingly challenging to reuse similar technologies to study human brains. Here we propose Adaptive Cell Tomography (ACTomography), a low-cost, high-throughput, high-efficacy tomography approach, based on adaptive targeting of individual cells suitable for human-brain scale modeling of single neurons to characterize their 3-D structures, statistical distributions, and extensible for other cellular features. Specifically, we established a platform to inject dyes into cortical neurons in surgical tissues of 18 patients with brain tumors or other conditions and 1 donated fresh postmortem brain. We collected 3-D images of 1746 cortical neurons, of which 852 neurons were subsequentially reconstructed to quantify their local dendritic morphology, and mapped to standard atlases both computationally and semantically. In our data, human neurons are more diverse across brain regions than by subject age or gender. The strong stereotypy within cohorts of brain regions allows generating a statistical tensor-field of neuron morphology to characterize 3-D anatomical modularity of a human brain.

## Introduction

Study of the human brain is one of the most significant endeavors of modern brain science (Purves, et al, 2019; Markram, et al, 2015; Luo, 2020). Despite the existence of many animal models to understand brains (e.g. Gouwens, et al, 2019; Peng, et al, 2021), it is still of utmost and profound importance to examine surrogates of human brain directly (DeFelipe, 2011; Farahany, et al, 2018). Acquiring neuron data from postmortem brains has been a powerful way to produce human brain map models (Hawrylycz, et al, 2012; Amunts, et al, 2013) and to discover new cell types (Boldog et al, 2018). Alternatively, screening of necessarily extracted brain tissues along a surgical path to remove brain tumors and/or other disease sites offers another opportunity to profile states of human brains. Previous efforts such as *in vitro* single cell characterization of human brains have started to generate 3-D morphology (Long, et al, 2017), electrophysiology and transcriptional profiles of single cortical L5 neurons from surgically extracted brain tissues (Kalmbach, et al, 2021). Over years, large datasets of single human neuron morphologies consisting of several hundred injected and traced human neurons were documented in frontal, temporal and occipital cortex (Elston, et al, 2001), anterior temporal lobe (Benavides-Piccione, et al, 2021), and CA1 (Benavides-Piccione, et al, 2020). Other relatively large-scale studies of human neuron morphology, until now, have been conducted using bulk staining such as Golgi (e.g., Anderson, et al, 2009; Hayes & Lewis, 1996; Warling, et al, 2021; Watson, et al, 2006). The field calls for a scalable platform for routine production of human neuron morphology with sampling and target-controlling capability.

Human *ex vivo* brain slices are often produced from surgically extracted brain tissues for collecting various physiological, morphology, and molecular attributes of individual cells. The *ex vivo* slices may also be sectioned from freshly dissected tissues in donated postmortem brains. Because of the requirement of minimal invasive surgical procedures, truncation of neurite structures is almost inevitable especially for long projecting excitatory neurons. Even for postmortem brains, damage of neurites is also hard to avoid during tissue extraction. On the other hand, in mammalian model animals such as mouse, it is known that quantification of individual cells’ structures at the whole brain scale is essential to study the complexity and diversity of cell types (e.g. Peng, et al, 2021). For human brains, although biopsy and autopsy tissues were also used (e.g. Benavides-Piccione, et al, 2020), it is an open challenge how to collect a sufficient amount of neuron morphology data to map and profile whole brains of individual neurons at the single cell resolution. The limited availability of genetic and viral labeling methods applicable to *ex vivo* human brain samples and the lack of computational analysis methods specifically developed for human neurons combine to form further challenges for the field.

We note that much of the existing effort in single cell characterization of human neurons is related to molecular, e.g. transcriptomic, analyses of cells (Lein, et al, 2017; Boldog, et al, 2018; Kalmbach, et al, 2021; Fang, et al, 2022). In general, the limited platform technologies to screen human neuron morphology also make it hard to connect single cell analyses of morphological and molecular attributes. Nonetheless, the recent advance of transcriptomic classification (Kalmbach, et al, 2021) and Patch-seq (Cadwell, et al, 2016; BICCN, 2021) characterization of single cells show the possibility to produce morphological classifier that could be used in a comparative analysis of these datasets. In addition, the increasingly available detailed maps of cell type distributions produced using spatial methods such as MERFISH (Xia, et al, 2019; Fang, et al, 2022), may help to make connections to neuron morphologies since there are very tight laminar distributions in different cortical areas. Therefore, it is becoming desirable to establish a scalable technical platform to routinely produce and analyze single human neuron morphometry, which would be iteratively refined to eventually facilitate classification of human neurons using large-scale conjugated analyses of molecular, physiological and morphological features.

As one of the first steps to address this massive challenge, this study proposes a low-cost, high-throughput method to screen human neurons at the human brain scale, based on *ex vivo* brain slices. Our method focuses on generating morphological profiles of individual neurons as a framework with potential extension to other cellular attributes. Here we describe this methodology and apply it to study both the diversity and stereotypy of human neurons in the morphological space. Our data suggests a hypothesis of morpho-gapping modularity of human brains at the single neuron resolution.

## Results

### Adaptive Cell Tomography and human *ex vivo* cell injection at whole brain scale

We developed an approach called Adaptive Cell Tomography, or ACTomography, to build 3-D neuron and distribution models from locally observed attributes of individual neurons (**Figure 1**). Here we limit the discussion to morphological attributes, or 3-D morphology, of human neurons. Our method begins with collection of small trunks of freshly dissected human brain tissues in craniotomy path to remove tumors/lesions in deeper brain regions (**Fig. 1A~C**), or tissues extracted from a postmortem brain. We produced thin, fixed brain slices (**Fig. 1D**), which allowed clear identification of neuron-somas using Differential Interference Contrast (DIC) imaging (**Fig. 1E, Methods**). We also used DIC imaging to guide the injection of dye (Lucifer yellow) into each of the selected somas (**Fig. 1E**). After the dye diffuses from a soma saturating the neurite region (**Suppl. Fig. 1**), we acquired a full 3D image stack of an individual neuron using laser scanning microscopy (2-photon) (**Fig. 1F** and **Methods**). We then traced a neuron’s morphology in 3-D to reconstruct its shape digitally (**Fig. 1G)**. This method was applied to as many neurons as possible on *ex vivo* brain slices, which were also mapped to the same spatial coordinate framework, to produce a large pool of digital morphology representations of individual neurons. Next, for every neuron we identified its clique of similar neurons (**Fig. 1H**), building a 3-D statistical neuron model based on its clique (**Fig. 1I**), retaining the single cell resolution at the whole brain scale. Statistical modeling of neurons, as shown later in this paper, also makes it easier to cluster brain regions.

**Figure 1.**
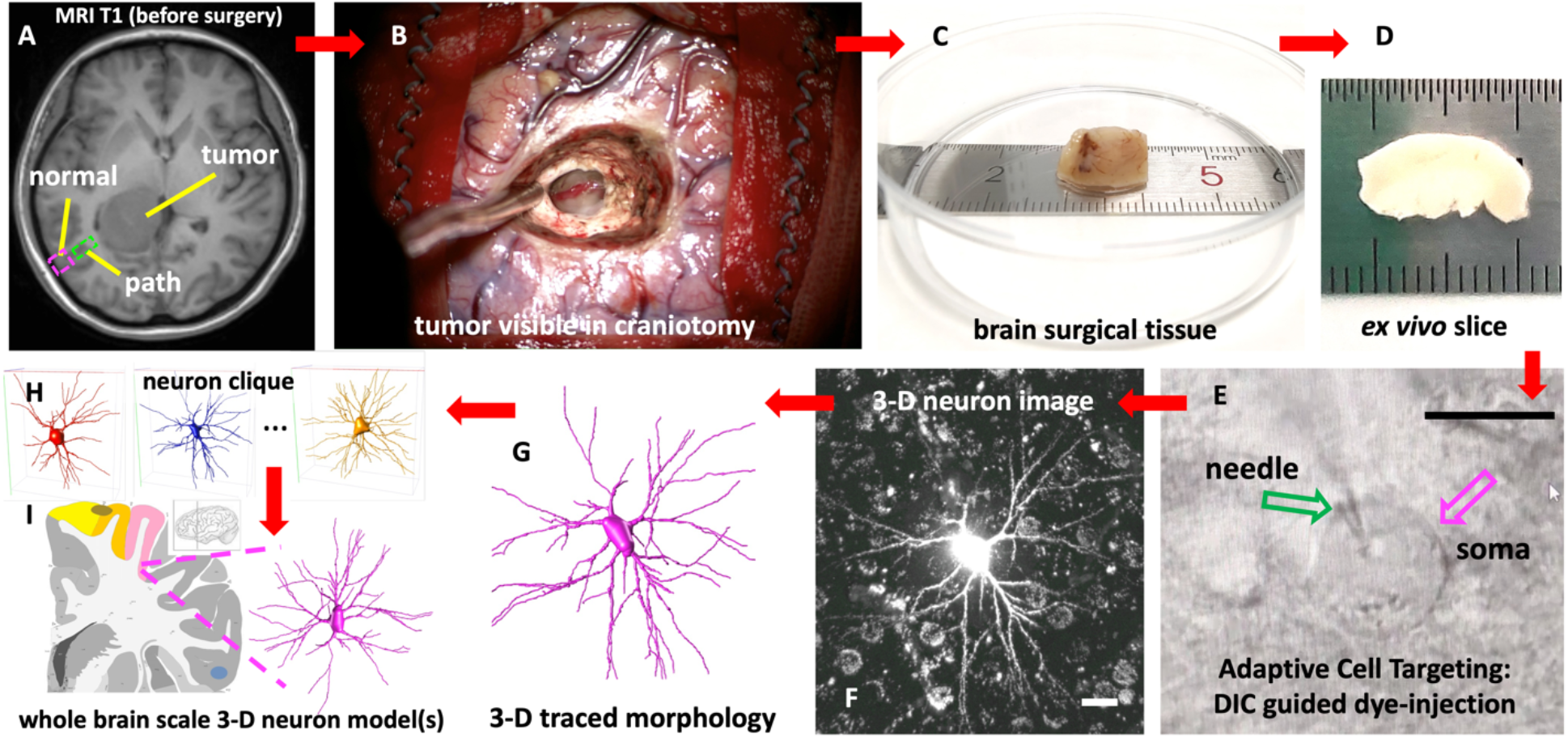
Schematic illustration of ACTomography for building neuron models in a number of brain regions at single cell resolution. Method is shown with surgically extracted brain tissues and is also applicable to postmortem brains. A. MRI T1 scan of a patient and schematic display of a small region (magenta) from which “normal” brain tissues would be extracted during surgery to remove the deep tumor/lesion along the “path” (green). B. Example of a tumor in surgery. C. Extracted human brain tissue, ~1cm^3^ in this case. D. 200μm thick ex vivo slice. E. DIC imaging of the brain slice. Scalebar=20μm. F. 3-D 2-photon 3-D imaging of an injected neuron (MIP is showed). Scalebar=20μm. G. 3-D traced morphology of the neuron in F. H. The clique of neurons that share similar features with G. I. Statistical analysis of a neuron-clique to build a 3-D neuron model that is also mapped to the approximate standard brain region.

ACTomography has two major components: Adaptive Cell Targeting (ACT) and single cell tomography. Because the choice of somas for dye-injection can be both selective (i.e. applied only to specific neurons or neuron-types) and non-specific (i.e. injection into every possible type of neurons visible in DIC imaging), our approach is capable for targeting and profiling a wide-range of neurons *adaptively*, limited only by the availability of human *ex vivo* brain slices. Previously, we succeeded in developing ACT for both *in vivo* single cell electrophysiology (Long, et al, 2015) and 3-D morphology reconstruction of large human neurons of *ex vivo* brain slices (Long, et al, 2017) as well as electroporation based 3-D tracing of *in vivo* mouse neurons (Long, et al, 2015). Therefore, a key step to make ACT work for ACTomography is to sample a large enough number of neurons independently to increase the success rate in reconstructing 3-D models of neurons, based on limited and often truncated neurites in human brain slices. As our previous experimental protocol for ACT-based dye-injection into human neurons was time-consuming, here we developed an inexpensive, high-throughput method to increase the sampling rate of neurons to improve the usage efficiency of human-tissue, while at the same time increase the availability of human brain specimens especially in different brain regions.

We thus established a whole brain scale workflow to collect human *ex vivo* cell injection images. Human *ex vivo* brain slices (thickness = 200μm in most cases) that contain “normal” neurons were produced for 8 males and 10 females, who suffered from severe tumors such as glioblastoma or other neurological conditions, but survived after brain surgeries, and the donated postmortem brain of 1 male died of glioblastoma (**Suppl. Table 1**; **Methods**). To maximize the potential to use brain tissues from subjects available only at different geographical regions, we introduced a fixation-and-injection approach so that *ex vivo* tissues could be prepared and shipped out to our processing center one thousand kilometers away without severe restriction of the freshness of brain tissues at the time of injection (**Methods**). For each neuron, we controlled the total dye-injection and diffusion time to be around 18.6±3.4 minutes (**Suppl. Fig. 1**), which could be further parallelized, followed by two-photon imaging to acquire a 3-D image stack of the neuron quickly (*t* = 98.1±23.7s). In this way, we acquired totally 3-D image stacks of 1746 neurons (**Suppl. Fig. 2** and **3**) out of 3231 injected neurons in 8 major non-overlapping brain regions and 2 overlapping regions (semantically in **Fig. 2A** and more precisely in **Figure 3**). Currently, this dataset is among the largest human single-neuron archives that has been produced. The aggregated production rate of both efficacy in image acquisition over cell injection (54.0% on average, within the range 25%~80% depending on the actual human tissue samples) (**Methods**) and neuron image production speed (~20 minutes/neuron) in *ex vivo* slices was at least 20 times more efficient than our previous cell-injection attempt (e.g. Long, et al, 2017).

**Figure 2.**
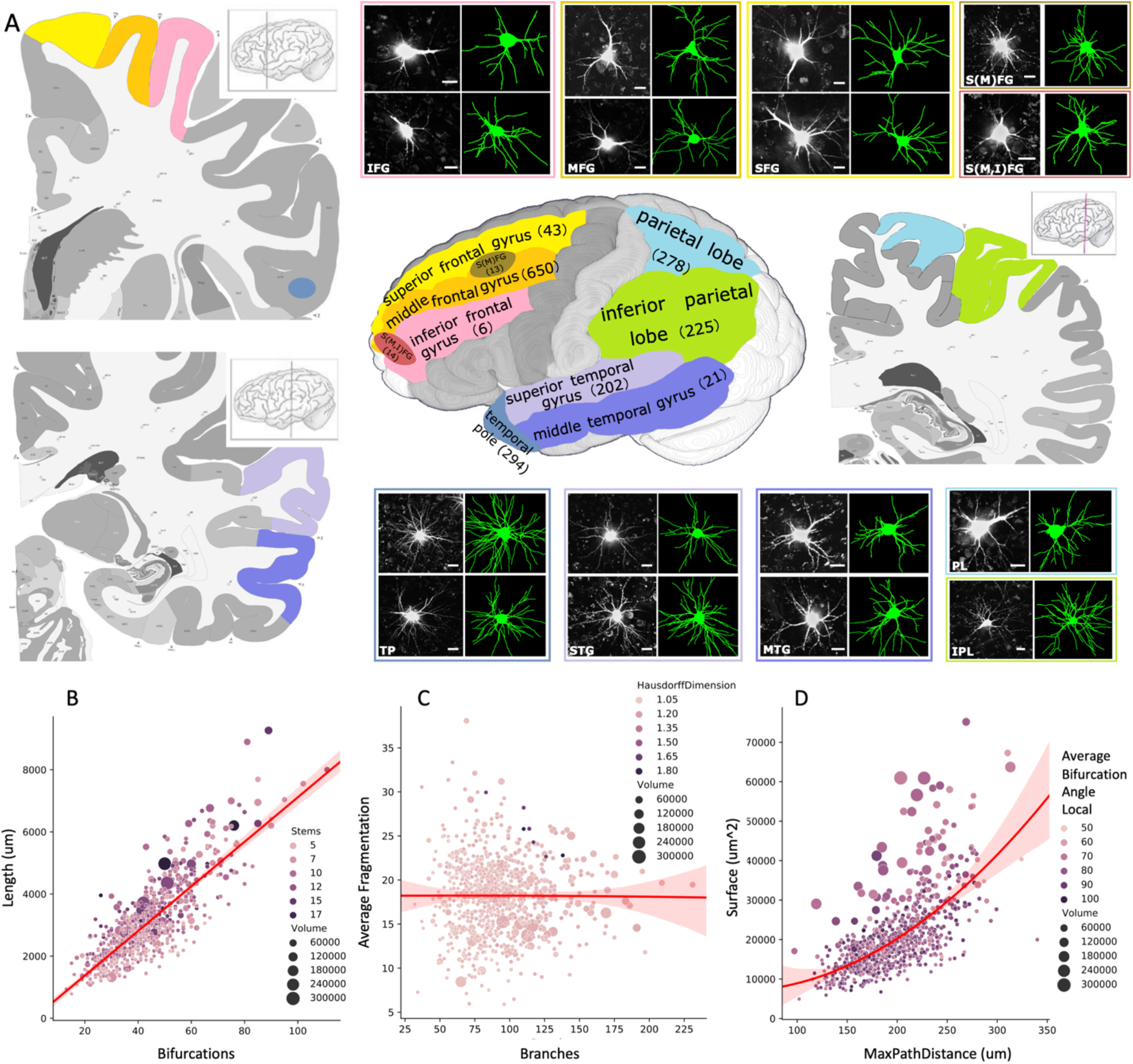
Semantic mapping of 1746 3-D injected, imaged, and 852 traced human neurons in 8 exclusive brain regions (irregular-shape) and 2 overlapping regions (oval) at whole brain scale (abbreviations in **Suppl. Table 1**), along with feature analysis of traced neurons. A. Sematic mapping scheme. Each brain region is shown in a different color along with the number of imaged neurons in it. Images (maximum intensity projection) and morphology (green) of one or two representative neurons are also shown for each region (some images are zoomed in for clear visualization; scalebar = 20μm). Note the actual locations of brain tissues extracted in this study as more precisely illustrated in **Fig. 3**, were much smaller than the color-labelled areas here. The brain template is based on Ding et al (2016). B. 4-D feature relationship among neuron bifurcation, length, number of stems, and volume of 852 neurons. C. 4-D feature relationship among neuron branching number, average fragmentation score, Hausdorff dimension, and volume. C. 4-D feature relationship among neuron maximal path distance, surface, average bifurcation angle (local), and volume. Optimal polynomial fitting curves (red) are also shown for B, C, D.

**Figure 3.**
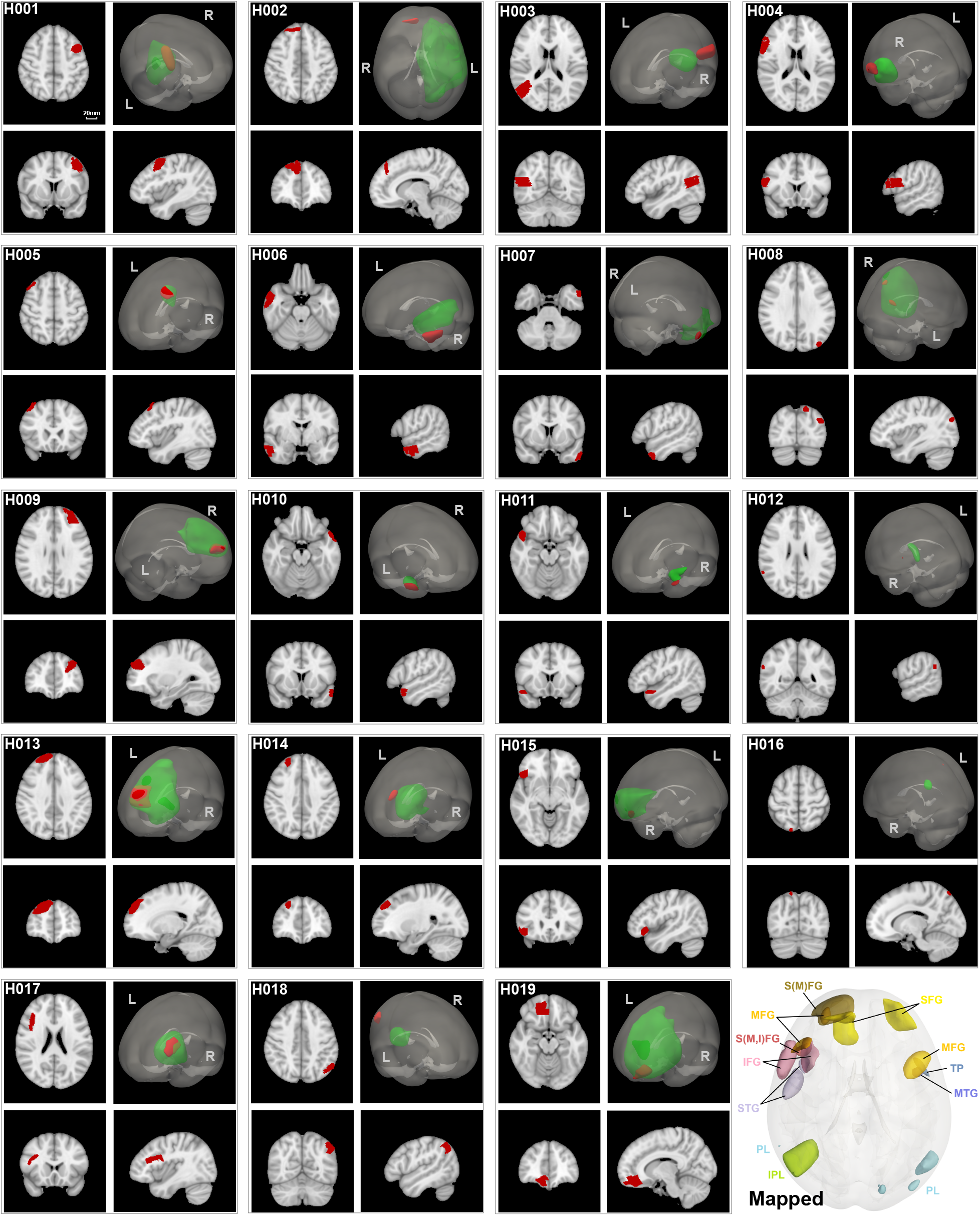
Whole brain computational mapping of tissue-extraction regions of human ex vivo brain slices, based on registration of post-operational MRI scans (for surgical tissues) or the last MRI scans (for postmortem brains). For each panel H001 ~ H019, the cross-sectional view shows the envelope of brain tissue extraction region (red), 3-D registered to a standard brain coordinate template MNI152. Scalebar = 20mm and the same for all cross-sectional panels. The 3-D views show both disease sites (green) and brain tissue extraction regions (red). Bottom-right shows all brain tissue extraction regions, each in a different color, mapped and overlaid on MNI152.

In the dye-injection step, as we have not restricted to specific neuron-types except the soma-visibility in DIC imaging, our current collection more likely contains large cells like pyramidal neurons in L3 and L5. To validate the acquired images recapitulate the “true” neuron morphology faithfully, we also used our approach to inject 491 mouse neurons in freshly dissected whole mouse brains. The efficacy of image acquisition over cell injection was 83.1%, reasonably higher than the upper range of efficacy in our human cell injection for good *ex vivo* tissues, due to the overall better condition to extract and utilize mouse brain tissues. The resultant 3-D image datasets of 408 mouse neurons were also compared to Cre labelled and fMOST imaged mouse neurons (Peng, et al, 2021) of the same types, as exemplified in caudoputamen neurons (**Suppl. Fig. 4**).

### Whole brain mapping of 3-D imaged and traced human neurons

We mapped the locations of extracted brain tissues and the accordingly acquired neuronal images to standard brain atlases, both computationally (**Figure 3** and **Methods**) and semantically (**Fig. 2A**). Of note, the actual brain tissues extracted for this study occupied only much smaller regions (**Fig. 3**) than the semantically depicted cortical regions (**Fig. 2A**). This whole brain mapping paradigm can assemble precious morphological, distributional, and associational information of single neurons.

We traced the 3-D morphology of 852 neurons that arborize heavily in dendrites (**Methods**). The traceable neurites are abundant during careful visual inspection with adjusted contrast of the neuronal images (**Fig. 2**). The five largest brain regions containing these traced neurons are middle frontal gyrus (MFG) (*n*=304), temporal pole (TP) (*n*=174), superior temporal gyrus (STG) (*n*=131), inferior parietal lobule (IPL) (*n*=112), and parietal lobe (PL) (*n*=91). We calculated 22 global morphological features (**Figure 2B~D, Methods**), which are independent of the orientations of neurons. Distributions of exemplar global morpho-features are also shown in **Suppl. Figs. 5 and 6**. Complex relationships among these features could be modeled using various data fitting methods, but also exhibit clear redundancy (**Fig. 2B-D)**. Therefore, the feature space was further reduced to 6 dimensions {#Stems, #Branches, Length, Average Bifurcation Angle - Local, Average Bifurcation Angle - Remote, Hausdorff Dimension}. The two pairs of features, {#Branches, Length} and {Average Bifurcation Angle - Local, Average Bifurcation Angle - Remote}, show well expected correlation among themselves (**Figure 4** – pair-plot) that also indicates neuron morphology in this study is consistent as previously documented human neuron morphology at NeuroMorpho.Org (Akram, et al, 2018). The distribution of #Stems and Hausdorff Dimension features is also consistent with previous documented human neurons. Overall, these two pairs of features and two remaining features #Stems and Hausdorff Dimension form a compact subspace to characterize the feature space of traced neurons in these five brains regions (**Fig. 4**), also in all 10 brain regions (**Suppl. Fig. 7**).

**Figure 4.**
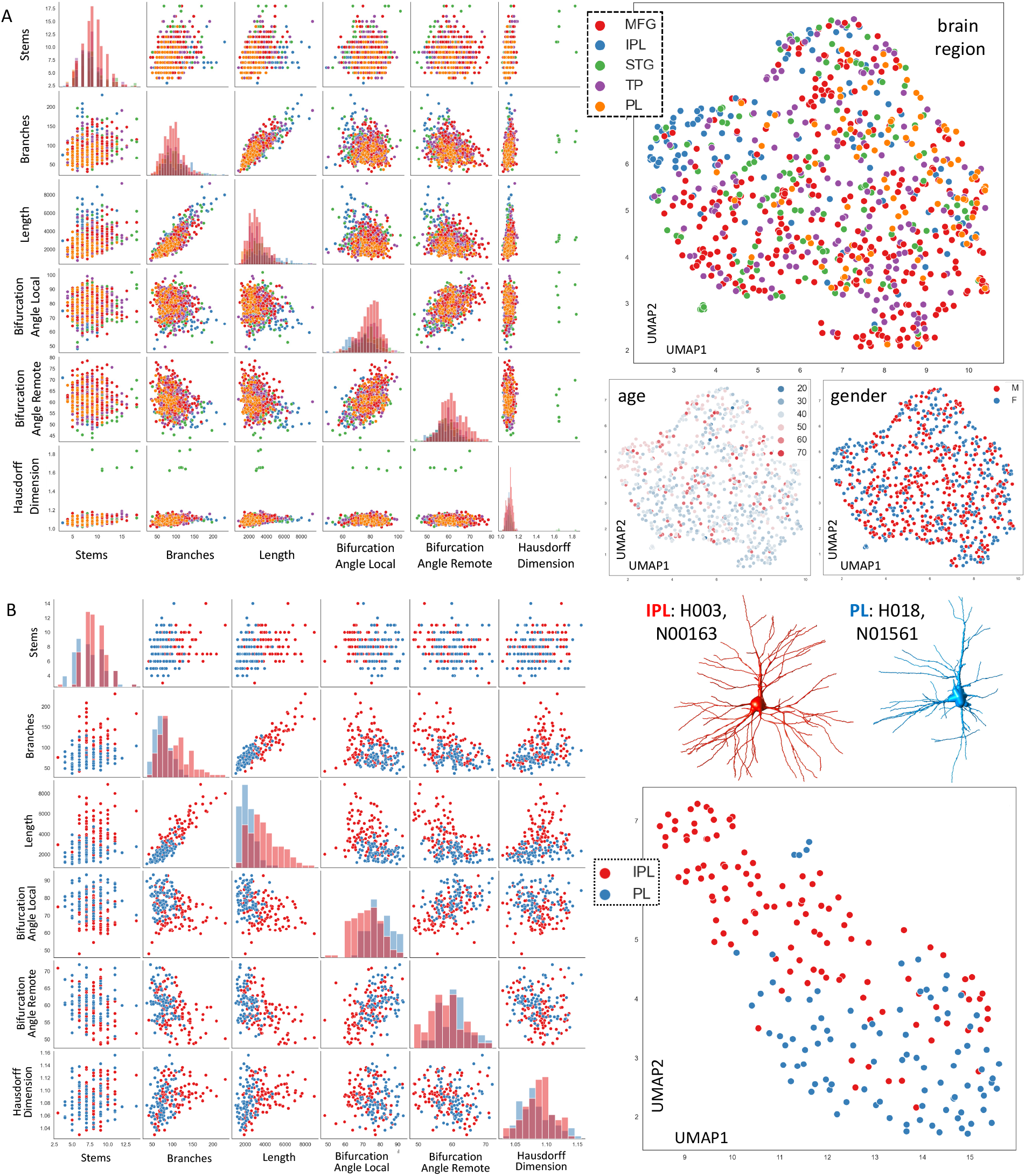
Analysis of global morphological features indicates greater diversity of human neurons across brain regions than age or gender of subjects and the stereotypy of neuron distribution patterns. A. Analysis of the 5 brain regions with most reconstructed neurons. Pair-plot of feature distributions (left): 6 key global morphological features reduced from the original 22-dimensional space. UMAP-plots (right): the feature space was further reduced using UMAP along with color-codes indicating three different groupings by brain region, age, and gender. B. Detailed analysis of two brain regions, PL and IPL, similar to A. Two examples of 3-D traced neurons for each region are shown, along with patient ID and neuron ID.

### Diversity and stereotypy of human neuron dendritic arbors

We analyzed dendritic arbors of all traced neurons to study neuron stereotypy, which is closely related to the diversity of neurons’ morphology with respect to different groupings of our data. We first considered the orientation-independent morphological features. A Uniform Manifold Approximation and Projection (UMAP) analysis indicates IPL, STG, PL and MFG neurons form a visible gradient of their feature distribution, while TP neurons spread out more in the feature space (**Fig. 4A** – UMAP-plots). In clear contrast, the age and gender of the human subjects do not differentiate these neurons (**Fig. 4A**). Thus these neurons are more diverse across brain regions than the age or gender. UMAP-analysis also indicates that neurons’ 3-D morphology is not correlated with human subjects’ age and gender, and is more stereotyped within a specific brain region than across different brain regions (**Fig. 4A**). Particularly, the two neighboring brain regions IPL and PL have distinctively conserved neuron morpho-features, as indicated by all individual key features (**Fig. 4B** – pair-plot) and the UMAP analysis (**Fig. 4B** – UMAP).

Although the 22-dimensional morpho-features did not yet produce substantial overall differences between neurons with varying orientations, we further validated the interareal diversity of neurons by examining neurons that share similar orientations in the raw images. We classified 3-D traced neurons into three pools, Pool-A, Pool-B, and Pool-T. Pool-A contains neurons that have well aligned and visible apical dendrites in the XY imaging plane (**Suppl. Fig. 8A, Suppl. Fig. 9** and **Suppl. Fig. 2**). Neurons in Pool-B have their apical dendrite oriented axially along the Z axis, i.e. orthogonal to the XY imaging plane, so that basal dendrites are mostly visible in the raw images (**Suppl. Fig. 8B, Suppl. Fig. 9** and **Suppl. Fig. 3**). Cortical neurons in Pool-A and Pool-B also correspond to the conventionally named “Coronal” and “Horizontal” views in previous studies (private communication, Ruth Benavides-Piccione). Pool-T contains all neurons with a tilted orientation in raw images. When subgroups of neurons sharing similar orientations (**Suppl. Fig. 8**) were compared with the situation that all neurons were analyzed together (**Fig. 4, Suppl. Fig. 7**), we observed similar continuum distribution and/or separation patterns, providing additional data to show that human neurons have greater diversity across brain regions than across gender or age.

We also considered other types of neuronal features in the orientation-classified analysis. We clustered the Sholl analysis (Sholl, 1953) profiles of neurons that share similar orientations (**Figure 5**). It is clear that most neurons innervate heavily around 50μm from somata, while some subgroups arborize more dominantly around 150μm from somata. Pool-A and Pool-B neurons form 2 and 3 major subgroups, respectively. They have evidently lower entropy values than brain-region based groupings (using IPL, PL, STG, TP, and MFG that contain most traced neurons) or random subgroupings (Figure 5 - bar chart), implying the neuron-arbors do form clusters in the space of Sholl-analysis profiles. However, the gain of the normalized mutual information (NMI) between these subgroups and the 5 brain regions as examined (**Fig. 2** and **3**, and **Suppl. Table 1**) is 3~4 times greater than the NMI scores between random subgrouping and the brain-regions groupings (**Fig. 5** bottom-left bar-chart). This shows that the neuron classes that arborize in varying density correlate with the cortical regions we examined, consistent with observation in **Fig. 4** based on global morphological features. The joint pattern of (a) clear clustering in the feature space and (b) the correlation with brain region distribution indicates that different types of neuron dendritic arbors in our data may spatially interlace with brain regions, potentially form a manifold across human cortical regions.

**Figure 5.**
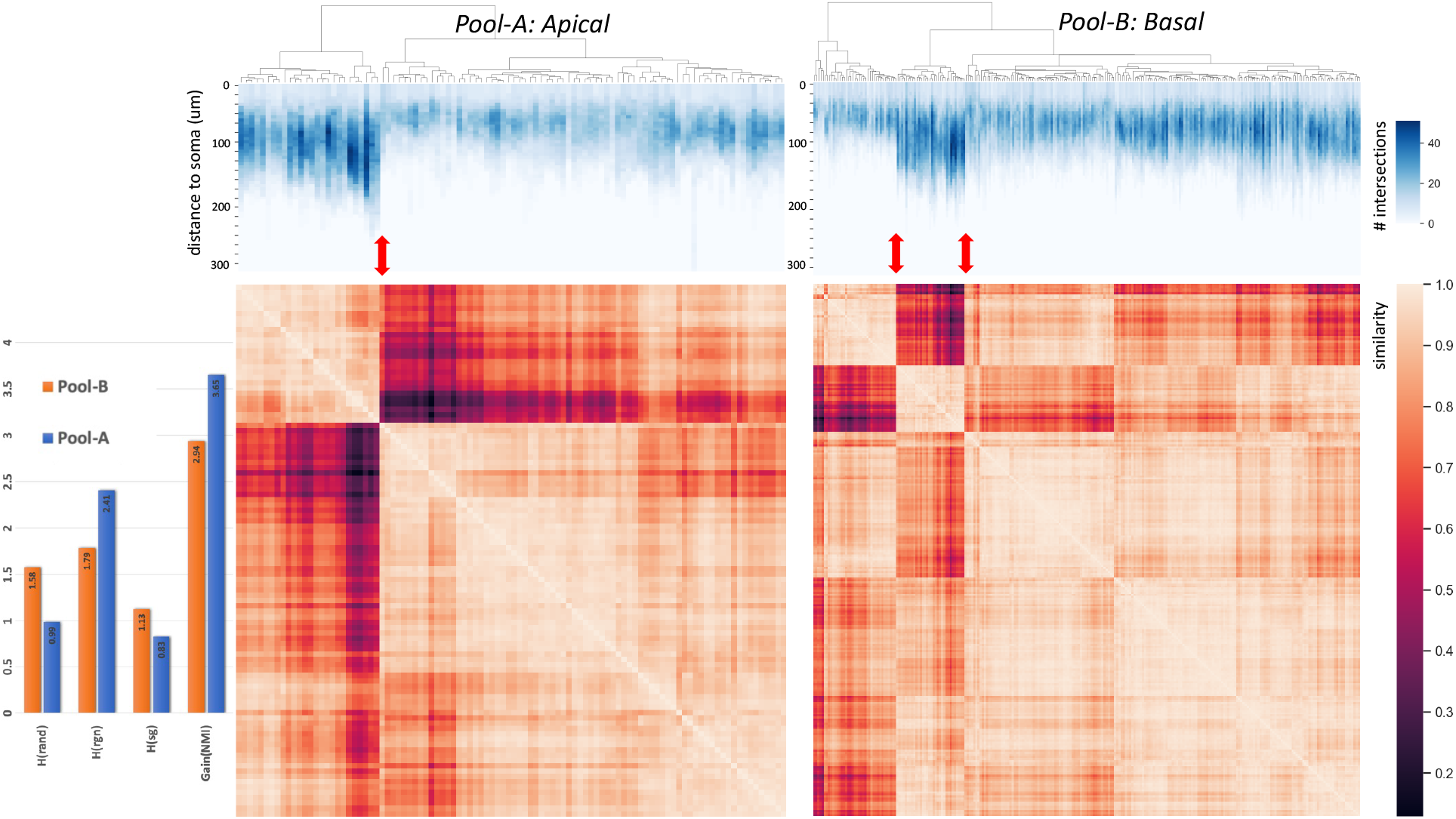
Orientation-classified analysis of human neuron distribution. The top-panel shows the Sholl analysis profiles, i.e. the number of branches of a neuron at specific distance to its soma. Neurons in Pool-A and Pool-B are separately clustered based on cosine similarity. For clear visualization of the generalizable similarity matrices, the sorted Pearson correlation (similarity) matrices matching the orders of clustered neurons at the top-panel are also aligned at the bottom panel. Pool-A and Pool-B neurons can be sub-grouped into 2 and 3 clusters, respectively, as indicated by red double-arrows. The entropy values of such sub-groupings (H(sg)), random sub-grouping (H(rand)), as well as the brain region based sub-grouping (H(rgn)) are shown in the left-side bar chart. The gains of the normalized mutual information scores are also displayed.

### 3-D neuron models and distribution models at the human brain scale

The strong morphological stereotypy of neurons allows us to use feature search, similar to BlastNeuron (Wan, et al, 2015), to detect N nearest neighbors of any target neuron, ideally also within each pool, to form a clique of similar neurons (**Figure 1H**). This method filters out morphologically dissimilar neurons that may be close in global feature space due to orientation. For each neuron-clique, it is reasonable to assume individual neurons are more likely from the same neuron type and share similar neuronal features. It then allows building a statistical neuron feature model that could generate new simulated neuron structures in 3-D (**Fig. 1I**). Specifically, using the 22-dimensional morphological features as an example, we detected the top-5 nearest neighbors in the feature space, and computed the mean and median feature vectors, as well as their variation feature vectors. This led to a 66-dimensional mean feature vector and a 66-dimensional variation vector of each neuron. We also estimated the overall mean and variation feature vectors for all traced neurons as the global background features, which were used to standardize the individual mean and variation features of single neurons. Optionally, we further normalized the mean vector using the respective variation, so that individual feature dimensions could also be compared with each other.

We examined the distribution and their associations among morphological features by removing global background. We found that morphological features in different brain regions could be well modeled by co-varying patterns. For instance, for three anatomically nearby brain regions inferior/middle/superior frontal gyri, the mean-features in SFG tend to co-vary linearly with those of MFG (negative correlation) and IFG (positive correlation), but are relatively independent of the variation-features of these regions in a 4-dimensional correlation analysis (**Figure 6A** - top-left pane). However, the variation-features of these brain regions do not exhibit clear correlation among themselves, and are also independent of the mean-features (**Fig. 6A** - middle-left pane). When the variation-features were used to further normalize the mean-features, we observed a strong condensation of features to the range of 0.75 to 1.5, while the entire distribution could be modeled by a 2^nd^-order polynomial (**Fig. 6A** - bottom-left pane). Very differently, for another pair of nearby brain regions, parietal lobe and inferior parietal lobule, their mean-features cannot be modeled linearly (**Fig. 6A** - top-right pane), while their variation-features exhibit weak negative correlation (**Fig. 6A** - middle-right pane). However, the variation-normalized mean features are also condensed to a narrow window of 0.8 and 2 while the entire normalized distribution can be approximated by a 2^nd^-order polynomial (**Fig. 6A** - bottom-right pane).

**Figure 6.**
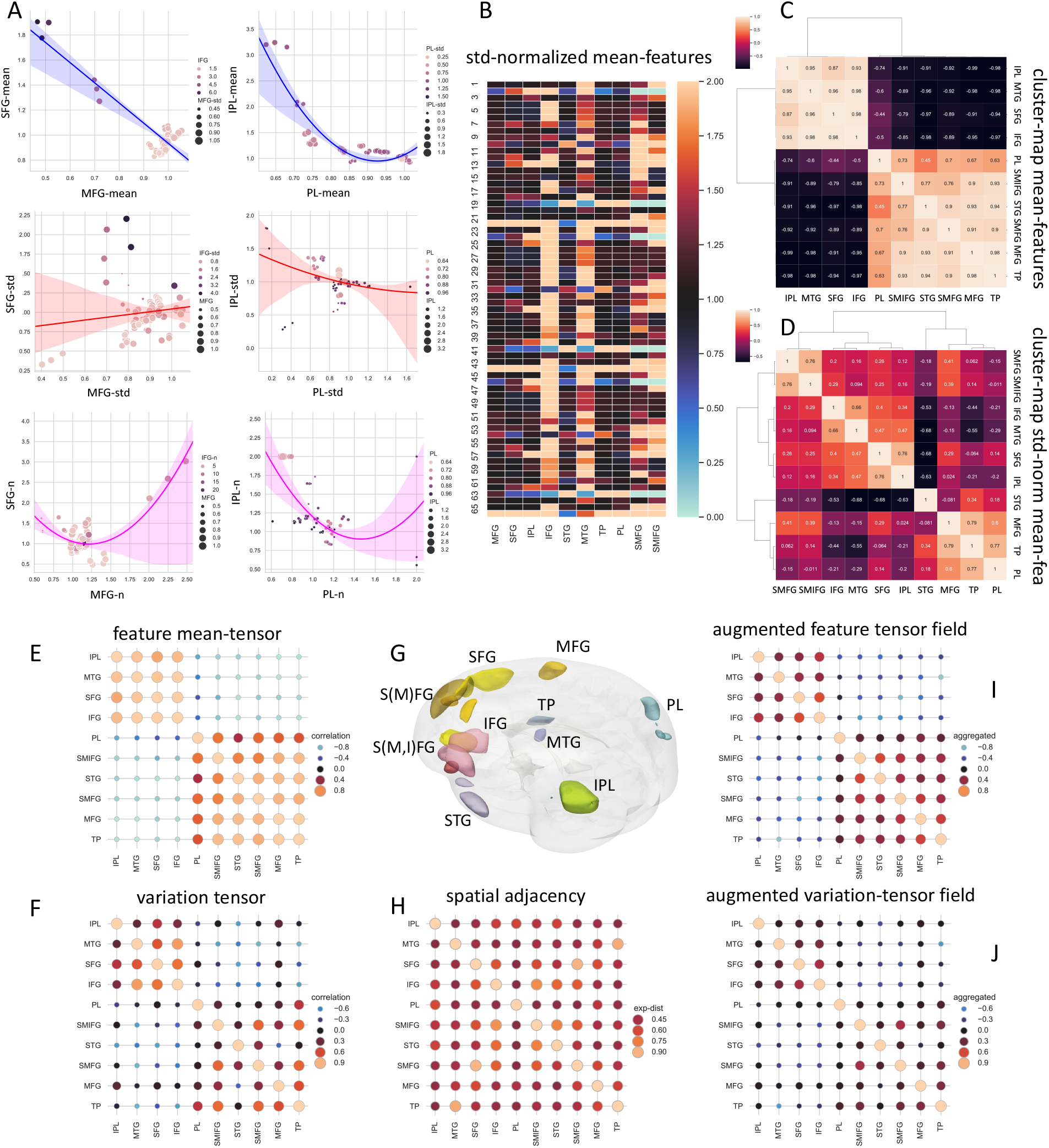
Generation of the brain-region-wise neuron-feature tensor field and the respective variation tensor field at the single cell resolution. A. Brain-region-wise multi-dimensional correlation analysis of mean feature vectors (top panes), variation features (middle panes), and the variation-normalized features (bottom panes). ‘-std’: standard deviation. ‘-n’: normalized by standard-deviation. Note both color and size of the data points are used to indicate additional dimensions other than x- and y-axes in the analysis. Optimal regression curves are also shown, with translucent regions indicating 95% confidence intervals of regressions. B. Variation-normalized 66-dimension mean-feature map of all brain regions. Feature value-range was clipped at 2 for clear visualization. C. Bi-clustering map of mean-features; the cosine-similarity is shown in each bin. D. Bi-clustering map of variation-normalized mean-features in B; the cosine-similarity is shown in each bin. E. The sorted mean-feature tensor; sorting order is the same as in C. F. The sorted variation-feature tensor; sorting order is the same as in C. G. 3-D visualized ex vivo brain regions of this study; color scheme: same as in **Figures 2 and 3**. H. Spatial adjacency map of brain regions in this study. Euclidean distances, d, between pair-wise representative medoids of extracted and 3-D registered brain tissues were computed; spatial adjacency = exp(-d). I. Spatial-adjacency augmented mean-feature tensor field. J. Spatial-adjacency augmented variation-feature tensor field.

We used mean-features and variation-features to investigate the distribution of single neurons in different brain regions. The variation-normalized mean-features characterize different brain regions clearly (**Fig. 6B**). Bi-clustering the similarity among these region-wise feature vectors shows details of neuron distribution patterns in brain anatomical regions. Indeed, we identified two remarkably clear cohorts of brain regions {IPL, MTG, SFG, IFG} and {PL, S(M,I)FG, STG, S(M)FG, MFG, TP} that have different morphological mean-feature vectors (**Fig. 6C**), while the variation-normalized mean-features are bi-clustered in a greater hierarchy across brain regions (**Fig. 6D**). In both cases, PL versus IPL features are evidently far away from each other. We also noticed that in both cases S(M,I)FG and S(M)FG features are close to MFG, which indicates that the respective traced-neurons might be actually more adjacent to or in the MFG region. In addition, in both cases, SFG/IFG features always differ from MFG features, consistent with the negative correlation observed in **Fig. 6A** (top-left pane). Together with the observed contrast between the PL/IPL pair, our data suggests a hypothesis of gapping modularity of neuron morphology distributed across nearby brain regions.

Therefore, for the first time we were able to further model the brain-scale interareal association of individual neurons’ morphology using tensors, which describe multi-region multi-dimensional statistics of neurons across the brain (**Fig. 6E~J**). Particularly, we generated the mean-tensor (**Fig. 6E**) and variation-tensor (**Fig. 6F**) of the brain regions involved in this study, based on the two cohorts of anatomical regions identified in **Fig. 6C**. These tensors demonstrate clear modules of the organization of brain anatomy, indicating the dendritic features of neurons are strongly conserved within each cohort, but interlace across brain areas. Within each cohort, the dendritic features have different levels of variations, which could be further modeled by using additional combinations of features in future work. Considering the 3-D spatial locations of the extracted brain tissues (**Fig. 6G**), we also computed a spatial adjacency map of these brain regions (**Fig. 6H**). We aggregated the spatial closeness of brain regions and the feature tensors, and generated the spatially modulated morphological feature tensor field (**Fig. 6I**) and the respective variation tensor field (**Fig. 6J**). Both augmented tensor fields uncover the rich structures of dendrite profiles of individual human neurons that are embedded deeply in the neuron-wise distributions shown in **Figs. 4** and **5**. The strong stereotypy as exhibited by localized morpho-features also indicates statistical modeling of neurons’ 3-D shape and neuron distribution can be reasonably achieved by applying adaptive cell tomography at the human brain scale.

## Discussion

In this work we proposed ACTomography to acquire one of the largest known single neuron morphology datasets for human brain. Different from several recent successful single-cell morphology studies on model animals such as mouse (Winnubst, et al, 2019; Peng, et al, 2021) and monkey (Xu, et al, 2021), for human brains there lacked a suitable large-scale technical platform to conduct similar studies. The pioneering efforts of cell injection, imaging and morpho-analysis were promising (Elston, et al, 2001; Benavides-Piccione, et al, 2020; Benavides-Piccione, et al, 2021; Berg, et al, 2021; Hrvoj-Mihic, et al, 2017; Jacobs, et al, 2014; Long, et al, 2015; Long, et al, 2017; Wang, et al, 2015), yet also had a limited scale for human brains. The primary goal of this paper is to provide a generalizable framework for setting up a high-throughput platform of human cell injection, imaging, neuron tracing, brain registration, pattern analysis, and modeling of neuron morphology and brain anatomy. Our approach is suitable for both brain tissues necessarily extracted in surgeries to remove deep brain tumors and lesions, or in donated postmortem brains. The rich information produced here shows that with this low-cost platform, we were able to achieve high-efficacy of successful neuron targeting and cell injection to produce reasonable 3-D dendritic morphology in fixed *ex vivo* brain tissues. The number of human neurons and mouse neurons in our experiments were large enough to provide statistical validation of our approach in comparison with previous studies of neuron labeling, imaging, and morphology analysis. The efficacy ratio in harvesting neuron data was also sufficiently high, which makes it desirable to boost the throughput further by automation and parallelizing soma detection, neuron-type identification, cell injection, and imaging.

We have used *ex vivo* brain tissues from 19 subjects covering 8 exclusive and 2 overlapping cortical regions, while at the same time used registration methods to spatially map these regions at the whole brain scale. This approach sets up a scalable framework to add more brain regions and cell types into this whole brain map of single cell morphologies. Currently, ACTomography allows us to adaptively select neurons whose somata are visible for injection. In cell selection, although here we focus on reporting injection and analysis of pyramidal neurons using DIC imaging, ACTomography is not limited by the imaging modality. First, various wide-field co-staining methods, such as DAPI labeling, or specific sparse labeling techniques, could be applied at the same time so that we may also inject dyes into a variety of potential cells, under fluorescent microscopes. Second, we could deploy an injection array or parallelize the injection to many loci simultaneously, and identify the successfully injected neurons in a *post hoc* way to triage specific neuron types. Our fixation-and-injection approach lowers the requirement of the freshness of brain tissues for injection. In turn, it maximizes the possibility to screen more neurons from different sources and conditions.

Our analyses provide a general assessment of the “background” dendritic model of neurons and their anatomical distribution, based on multi-dimensional correlation of morphological features and brain regions, of “normal” brain tissues. To map the diversity and stereotypy of neurons, we have used both semantic and computational brain mapping to pinpoint neurons’ anatomical locations. We found that dendritic features of human neurons in our data correlate with major cohorts of brain regions (cf. e.g., Benavides-Piccione, et al, 2006 for similar rodent finding and Jacobs, et al, 2001 for a human Golgi study). Such dendritic features do not exhibit strong correlation with gender or age of the subjects. The spatial distribution of brain regions can also be fused with the single-neuron-level dendritic feature maps as tensor fields to model and visualize the anatomical modularity and complexity implied by the diversity and stereotypy of morpho-features of neurons. This study provides an orthogonal, complementary paradigm to previous work on analyzing morphological property and distribution of mammalian neurons at the whole brain scale (e.g. Peng, et al, 2021). Rich information can be well anticipated when more data, especially metadata, become increasingly available in continuing efforts, such as the cortical layer, spiny, left and right hemispheres information of these neurons we would report in a sister-piece of this study, functional and behavioral data of the subjects, comparative data of “abnormal” cells in the near-lesion or in-lesion brain tissues, and many other brain regions not reported in this study.

In contrast to other model animals, one of the key challenges to study human neurons’ morphology at whole brain scale is about the completeness and fine details of neuron shape, especially arborizing patterns of long projecting axons and apical dendrites. An intrinsic limitation is due to the *ex vivo* nature of human brain research in this direction. For single-cell injection technologies, an obvious improvement is to optimize tracer (e.g. dye) diffusion along neurites. While we may switch tracers and also keep upgrading tracer injection techniques, we note that one strength of ACTomography is the fast and effective accumulation of a number of locally visualized human neurites in precisely pinpointed anatomical regions. The strong stereotypy of intra-region neuron features suggests two approaches to obtain more complete shape information of individual neurons. First, for an experimentally observed target neuron, we may aggregate local information of its neurites in a mean model along with estimated variations of morphological dimensions, as shown in **Fig. 6**, and therefore synthesize the missing structures by learning from existing features of nearby neurons. Second, we may generate new neurons by integrating morphology tensor models over anatomical regions. When neurons’ orientations are aligned (e.g. **Fig. 5**), a silhouette of neurons’ shapes may be turned into a statistically sound representation to model fine details of neuron arbors. By putting together numerous pieces of neurites, a “shotgun fragment assembly” approach similar to whole genome sequencing algorithms (Weber and Myers, 1997) might be possible to reconstruct long projecting axons in human brains.

Finally, previous studies (e.g. Kalmbach, et al, 2021) and current large-scale brain science initiatives strongly suggest that the morphological profiling of human neurons at multiple brain regions should be combined with molecular and physiological profiling of individual neurons. ACTomography is generalizable to incorporate some of those experimental components. We envision a possible combination could be spatial genomics or *in situ* transcriptomics (Alon, et al, 2021) for the *ex vivo* brain slices. As we have successfully mapped various morphological tensors extracted from *ex vivo* samples over the standardized brain anatomy, the same brain mapping strategy is highly applicable to fuse such multi-modality single-neuron resolution data at the whole brain scale. In addition, one may also use genetic tools such as enhancer AAVs (Graybuck, et al, 2021) to pre-label specific cell types genetically in cultured human tissues (Bakken, et al, 2021; Kim, et al, 2022). Incorporating such a pre-labeling method could be highly beneficial to the adaptive cell targeting step of ACTomography, or the *post hoc* triage of neuron types as discussed above.

## Acknowledgement

We thank Jie Xue, Lulu Yin, Yuanyuan Song for generation of neuron data, Brian Long, Songlin Ding, Ruth Benavides-Piccione, Javier DeFelipe, Rafael Yuste, Jonathan Ting, Linus Manubens-Gil, Yimin Wang, An Liu for various inspirations including discussions, comments, and suggestions to this line of work, Weiguo Tao for providing the micropipette puller, Yichen Guo, Zhixian Gao, Zhen Wu, Yang Zhang, Tao Sun, Xiong Xiao for support. This project was mainly supported by a Southeast University initiative for neuroscience awarded to Hanchuan Peng. Liwei Zhang was supported a Beijing Municipal Special Funds for Medical Research (Grant No. Jing Yi Yan 2018-7) and a grant from National Natural Science Foundation of China (Grant No. 81872048). Peng Zhang was supported by the Beijing Municipal Natural Science Foundation (Grant No. 7214214). The Southeast University team was also supported by a MOST (China) Brain Research Project, “Mammalian Whole Brain Mesoscopic Stereotaxic 3D Atlas” (2022ZD0205200, 2022ZD0205204) awarded to Lijuan Liu. Wei Xie was supported by a MOST (China) Brain Research Project (2021ZD0204001). Liya Ding was supported by Fundamental Research Funds for the Central Universities (2242022R10089). Giorgio Ascoli was supported by NIH grants R01NS39600, U01MH114829, RF1MH128693, and R01NS86082.

## Author Contribution

H.P. conceptualized this study and managed the entire project. L.Z. co-managed the project and provided brain tissue resource. H.P. invented the adaptive cell tomography method and related informatics techniques. H.P. analyzed the data with assistance of Z.Y., F.X., and S.G.. X.H. managed the web-lab experimental platform and co-lead the cell-injection and neuron image acquisition with S.G. and L.L. S.G. contributed to neuron analysis and developed data management with J.L., X.Y. and Jie Xue in Acknowledgement. N.J. and L.Z. provided brain tissues and all metadata of brain tissues, with help from Yi.W., Yujin W., Y.H., P.Z., W.W. Z.Y. and F.X. contributed to brain mapping and data analysis with assistance from T.L. J.L., X.Y. J.X., and J.R. performed cell injection and image acquisition. Lulu Yin, Yuanyuan Song in Acknowledgement and D.L., H.M. traced neurons, pooled neurons, and contributed to initial triage. L.L. co-supervised the team and contributed to critical data production of cell injection, imaging, and neuron tracing. E.L. and M.H. contributed to data acquisition and analysis strategies. G.A. contributed a comparison to existing human data in NeuroMorpho.Org and related metadata. L.D. computed Sholl analysis features and provided a tissue slicer. W.X. provided support for mouse injection comparison and analysis. H.P. wrote the manuscript with input from all coauthors.

## Methods

### Human specimens

A total of 19 human cases were enrolled in this study, with 18 cases obtained from surgical procedure in Beijing Tiantan Hospital and 1 case obtained from donated brain from Human Brain& Tissue Bank of National Clinical Research Centre for Neurological Disorders. All the cases were informed for consent and all procedures involving human tissue for research purposes were approved by the institutional review board of Beijing Tiantan Hospital.

Totally 8 male and 10 female patients between 18 and 71 years old underwent the surgical resection of primary tumor or similarly documented brain diseases for treatment purpose, with clear surgical indications. All the patients were confirmed by preoperative imaging that the brain tumor or lesion located in the deep brain structure or suspected for high-grade gliomas, which were indicated for surgical approaches path through the normal cortex following the neurosurgical principles. The surgical procedure cautiously preserved the normal cortex limited in the approached or in the edge of high-grade gliomas. Post-operative histological review confirmed the tumor/lesion pathology and function status were assessed by follow-up after patient discharge.

In the meantime, the case H002 was obtained from a donated patient died of glioblastoma, which extensively infiltrated in the left brain. The normal frontal lobe tissue was acquired from the right side following the protocols approved. Specifically, this patient received tumor resection on September 13, 2020, due to space-occupying lesions in the left frontal lobe. Postoperative pathology reported glioblastoma, and postoperative chemoradiotherapy and electric field therapy were given. The patient underwent the last MRI examination on Feb 23, 2022 and eventually died due to disease progression on Apr 22, 2022. After death, the brain tissues were donated to the Human Brain & Tissue Bank of National Clinical Research Centre for Neurological Disorders and dissected and preserved by qualified personnel. In this study, the right frontal lobe tissues of patients were obtained in strict accordance with ethical requirements, and the corresponding anatomical sites were judged by professionals and photographed for preservation.

### Brain tissue preparation and sectioning

The brain tissues were fixed, fluorescent labeled, and imaged. About 5-10 minutes (except two cases of large tissues, it was within 20 minutes) after the surgery, the extracted specimens were immersed in 4% paraformaldehyde (PFA) phosphate buffer (PB) solution (0.1 M, pH 7.4) for fixation. Specimens larger than 1 cm^3^ were cut into such a unit volume before fixation. Prior to sectioning, the tissues in PFA solution were stored on a shaking table at 4°C for ~24 h.

The fixed tissue blocks were then sectioned into slices before being labeled. The tissue blocks from the PFA solution were taken out to remove blood clots and meninges on the surface, followed by rinsing three times with double distilled water. It was then put in a dry petri dish, where water on the surface was removed using absorbent paper. Then we poured 3% agarose gel (42°C) into the Petri dish until the tissue is completely submerged, while removing the bubbles around the tissue. The excess agar blocks were removed after the agar had solidified.. Thereafter, we sectioned the tissue block into 200-300 μm thick slices (~80% 200μm thick) with vibratome (Leica vt1200s). During this procedure, the tissue block was fixed on the vibratome with 404 glue (LOCTITE) and immersed within 1×PBS. We cut the tissue longitudinally along the surface of the cerebral cortex, from gray matter to white matter. The slices were transferred to 5ml centrifuge tubes and stored in 4% PFA solution on the shaking table at 4°C for 3-5 days to be further fixed. The fixing brain slices were still placed.

### Fluorescent neuron labeling through cell injection

To distinguish individual neurons from surrounding cells, we labeled single neurons with Lucifer Yellow (YL) CH, Lithium Salt (ThermoFisher) through cell injection with micromanipulators. Injection was conducted under an upright differential interference (DIC) microscope (Scientifica 1000P) equipped with near infrared light (~900 nm) as light source and a water-immersion objective. As we found that Lucifer Yellow could be effectively injected into neurons of fixed tissue slices and label the neuron including dendritic tree and spines, and potentially also axons, under optimized conditions related to the specimen. The negatively charged YL was pumped into a neuron with the help of a micro electrophoretic current generator (WPI Microiontophoresis Current Programmer SYS-260).

The detailed procedure of cell injection is as following:

1. Brain slices were carefully transferred onto the sample stage of the DIC microscope and held confined with a silk wound copper ring tablet above it. The sample chamber was filled with sufficient amount of 1×PBS to keep brain slices immersed in PBS.
2. Pipettes needed for the cell injection were produced by a micropipette puller (Sutter instrument P-2000). Borosilicate glass capillaries (WPI 1B100-4, 1.0 mm o.d. × 0.58 i.d.) were employed to pull very fine pipette with a long needle tip (needle tip size 1-2 μm and needle tip angle ~10°). These pipettes also served as the microelectrodes of the microiontophoresis.
3. Pipettes filled with 4% LY PBS solution were mounted on a microelectrode holder (WPI, PE MEH710), which was fixed on a 3-D adjustable micromanipulor (Scientifica, MicroStar S-MST-1000-X). In this process, it is critical to have sufficient solvent and without bubbles in the in the pipette and holder. The negative electrode of the microiontophoresis device (WPI SYS-260) was connected to the microelectrode holder filled with LY solution. The positive electrode was connected to the tablet iron wire and immersed in PBS solution.
4. In DIC microscope, we used a 40x water immersion objective and a CCD to acquire the image. After the sample was placed, needle tips were moved to the field of vision of the microscope. Then, the objective and needle tips were moved down synchronously until the objective is well focused on the brain slice. The neuronal somas about 30-50 μm deep within the brain slice were selected for labeling. For a given neuron soma, the needle tip was first moved to the position directly above the soma and then slowly moved downward to approach the soma. The cell is punctured after the tip of the needle contacted the cell membrane. We know the needle penetrated the cell membrane if we could see an instantaneous rebounding of the cell membrane. The brain slices were imaged on live at DIC mode during this whole procedure.
5. After the needle penetrated into the cell, the microiontophoresis device was turned on, the diffusion of the LY in a neuronal soma as well as along dendrites can be observed via the wide field fluorescence imaging of the DIC microscope. The perfusion is continued with a negative current of 10 nA for 15-20 min until the branches were clearly visible. After the perfusion is finished and the microiontophoresis is turned off, we waited for another 5 minutes before pulling out the needle.

### Two-photon fluorescence imaging of neurons

We used a two-photon fluorescence imaging system (Bruker, Ultima Investigator) to obtain the fluorescence microscopic imaging of the labeled neurons. The imaging system was setup on an upright microscope with a single set of galvanometers providing standard and resonant scanning mode, 8 kHz x-axis scanning and high speed 30 fps for 2D 512×512 scan. For LY-stained neurons imaging, the fluorescence collection used was a filter module composed of 560 nm long pass dichroic beam splitter and an emission filter et525/70m-2p. The objective used was an apochromatic water immersion 25× two-photon objective (Nikon, CFI75 Apochromat 25XW MP1300) with a numerical aperture 1.1. Two-photon fluorescence excitation was provided by a Ti:sapphire femtosecond laser (Coherent Chameleon Discovery NX with TPC Laser System) generating widely tunable (680-1300 nm) femtosecond laser pulses (pulse duration <120 fs). The repetition frequency of the laser output pulses is 80 MHz, with an average output power of 2 W at 900 nm (used for LY labeled neurons imaging). The scan and image collection were controlled by the software of Prairie View. The brief two-photon imaging process is as following:

1. The brain slices were transferred to the glass slide and confined by a silk wound tablet. An appropriate amount of 1×PBS was dropped on the brain slices ensuring that the objective lens can be submerged in PBS when taking images.
2. The slides were put on the stage of two-photon fluorescence microscope. The wide field fluorescence imaging mode was used for the localization of target neurons.
3. The imaging system were switched to two-photon imaging mode after the cells were founded. Using Primitive view software, in the live scan mode, the XY and Z-axis could be adjusted to determine the imaging range of two-photon scan. In the stage control module, the upper and lower limits of the Z-axis scanning of cells and the scanning step length could be set. The imaging zoom in/out was adjusted according to the extension of neurons in the XY plane. After setting, Z-series scan could be started to get the image stack.

We defined the efficacy in image acquisition over cell injection as the ratio of the number of successfully imaged neurons with visible neuron arbors divided by the total number of attempted injections.

### Brain registration

All subjects involved in this study had complete MRI data. Based on operation records and the pre-operation and post-operation MRI data, we applied 3Dslicer to T1 images to define the loci of the extracted brain tissue. For instance, for H003, we used 3D Slicer to load DICOM format T1 imaging data, and then used the Segment Editor module of 3D slicer to produce the outline of the extracted brain tissue. Then, both T1 image and the segmentation were output as files (in nrrd format).

We used the Brain extraction Tool in FMRIB’s Software Library (FSL) to segment T1 images and exclude skull in images (fractional intensity threshold = 0.5). We used MNI152 template included in the ANTS software (Avants, et al, 2009) as the target brain for registration, with the ‘SyN’ elastic alignment method. The tumor and other lesion regions were manually defined in the T1-registrered images. When the automatic registration quality was not satisfactory, optionally we used 3D Slicer to define the locations of extracted brain tissues on the MNI152 template directly.

### Neuron Tracing, Neuron Feature and Data Analysis

All 3-D neuron morphologies (in swc format) were traced and reconstructed using Vaa3D (vaa3d.org; Peng, et al, 2010), an open-source and cross-platform system for visualization and annotation, by a group of annotators in a semi-automatic way. In particular, two modules, TeraFly (Bria, et al, 2016) and TeraVR (Wang, et al, 2019), which enable large-scale non-immersive and immersive neuron reconstruction, were used to trace these neurons. Potential reconstruction errors were identified and removed through iterative quality control and correction. Thereafter, the morphology skeletons were refined to the center of the fluorescent signals via mean-shift algorithm, followed by pruning and sorting to generate neat neuron reconstructions, i.e., a single tree without breaks, loops, or unexpected branching. The root node of a neuron tree was set to be its soma. When needed, the contrast of the image was adjusted during the reconstruction to trace very weak neurites. It also helped to better visualize distal neuron fibers that might be less stained by fluorescent dyes during the cell injection. Neuron orientations were manually determined using Vaa3D based on clearly oriented apical dendritic stem(s) or the basal dendrite profiles.

The 22-dimensional morphological features include the following {‘Nodes’, ‘SomaSurface’, ‘Stems’, ‘Bifurcations’, ‘Branches’, ‘Tips’, ‘OverallWidth’, ‘OverallHeight’, ‘OverallDepth’, ‘AverageDiameter’, ‘Length’, ‘Surface’, ‘Volume’, ‘MaxEuclideanDistance’, ‘MaxPathDistance’, ‘MaxBranchOrder’, ‘AverageContraction’, ‘AverageFragmentation’, ‘AverageParent-daughterRatio’, ‘AverageBifurcationAngleLocal’, ‘AverageBifurcationAngleRemote’, ‘HausdorffDimension’}, defined in L-Measure (Scorcioni, et al, 2008). We used the implementation in Vaa3D’s global_neuron_features plugin.

The Dendritic Field Area (DFA) (Oga et al., 2017) was calculated to benchmark the dendritic spread of the neurons. Accordingly, we projected each neuron reconstruction onto the x-y plane and calculated the area contained within the convex hull around the dendritic terminations. The calculation was performed separately for neuron reconstructions of Pool-A and Pool-B to take consideration of the neuron orientations.

Data analysis was done mostly in Python, with the Python Data Analysis Library (pandas) and Statistical data visualization library (seaborn).

## Supplementary Figures

**Supplementary Figure 1.**
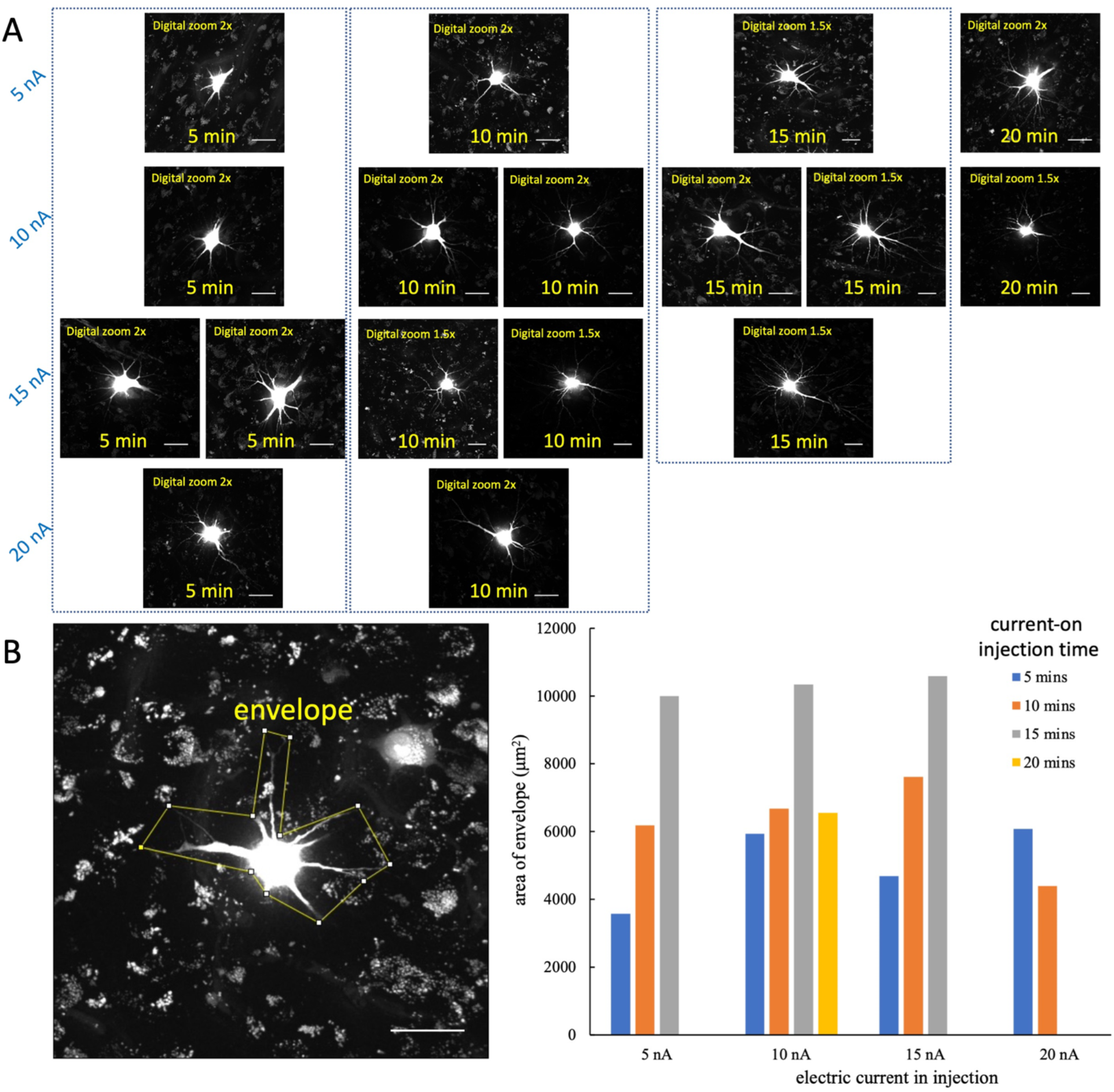
Optimization of dye-injection time and electric current for human neurons. For all the imaged human neurons in this study, the electric-current-on time for cell injection was 13.80±4.47 mins, followed by a current-off dye-diffusion time 4.76±1.30 mins. A. Examples of injected and imaged human neurons under different electric current and injection time conditions. Scalebars = 20μm. B. Example of an envelope of a neuron that contains all visible neurites. This envelope was determined by a technician carefully for each neuron during the setting-optimization. Scalebar = 20μm. C. Relationship between area of the envelope and two key parameters {electric current (nA), current-on time (minutes)}.

**Supplementary Figure 2.**
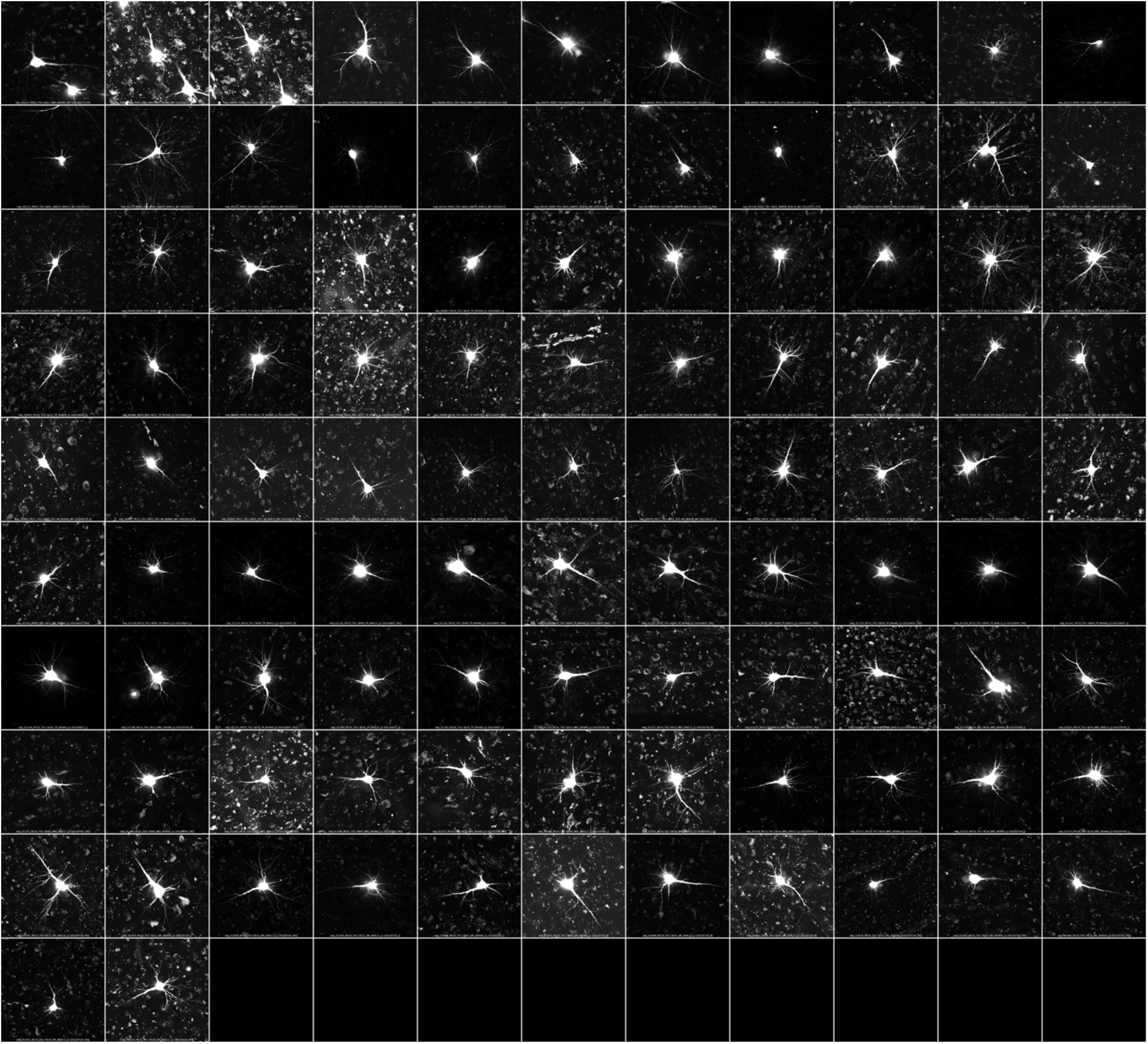
Apical dendrites of 101 human neurons. Zoom-in to see full details.

**Supplementary Figure 3.**
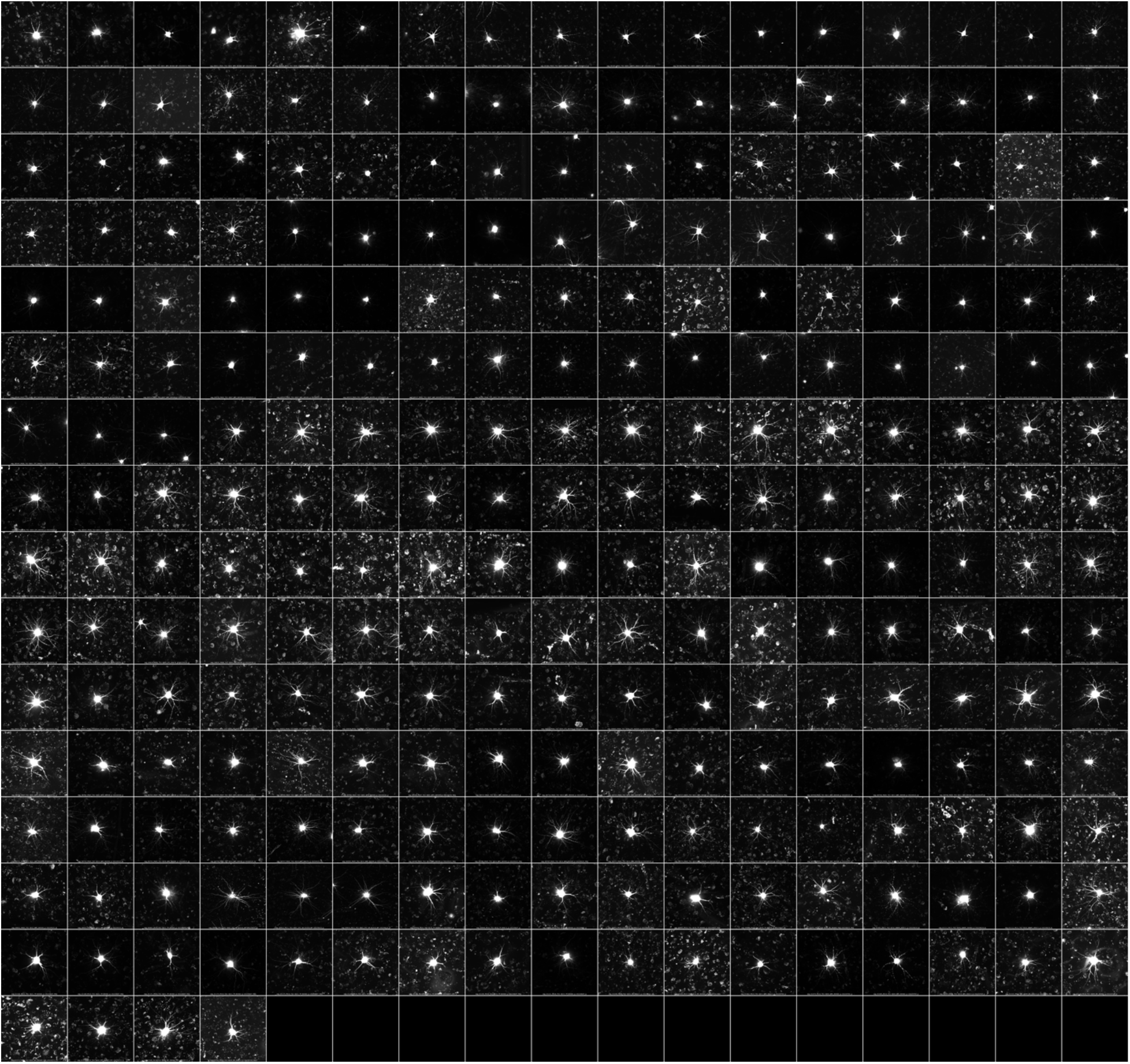
Basal dendrites of 259 human neurons. Zoom-in to see full details.

**Supplementary Figure 4.**
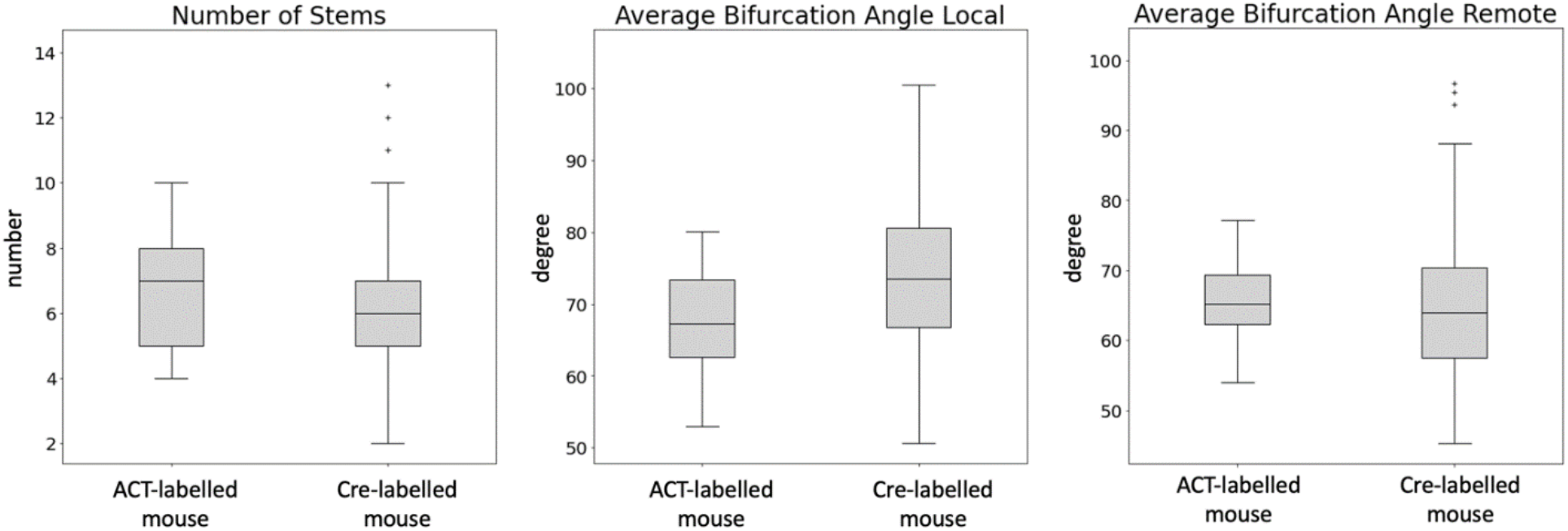
Comparison of three size-independent morphological features of ACT-injected mouse caudoputamen (CP) neurons (n=30) and Cre-labeled and fMOST imaged mouse CP neurons (n=311).

**Supplementary Figure 5.**
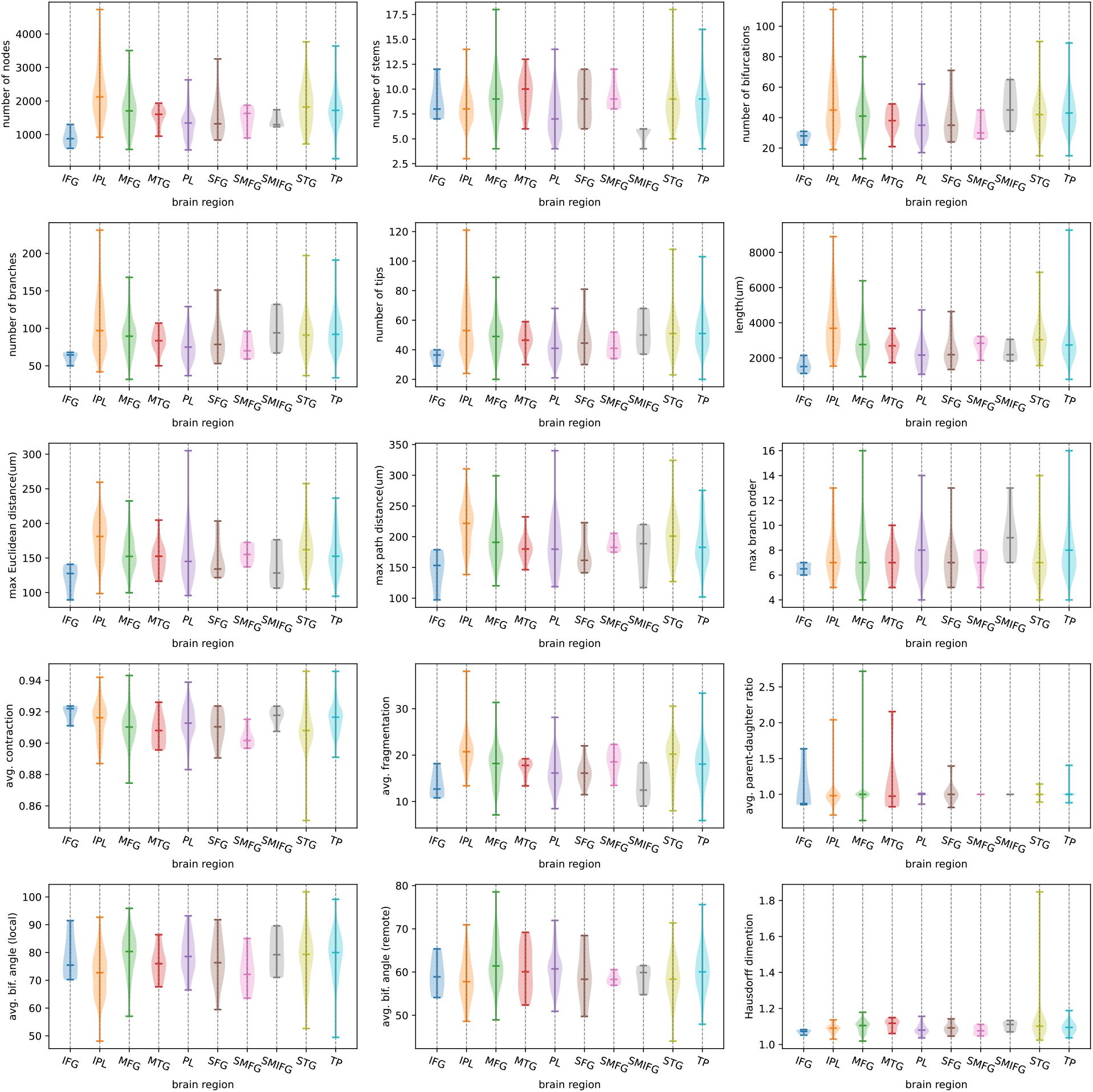
Exemplar global morphological features of each neuron reconstruction visualized as violin plots, grouped according to the brain region. Each violin set shows the distribution of the values from a particular group, with the median value indicated by the bar in the middle.

**Supplementary Figure 6.**
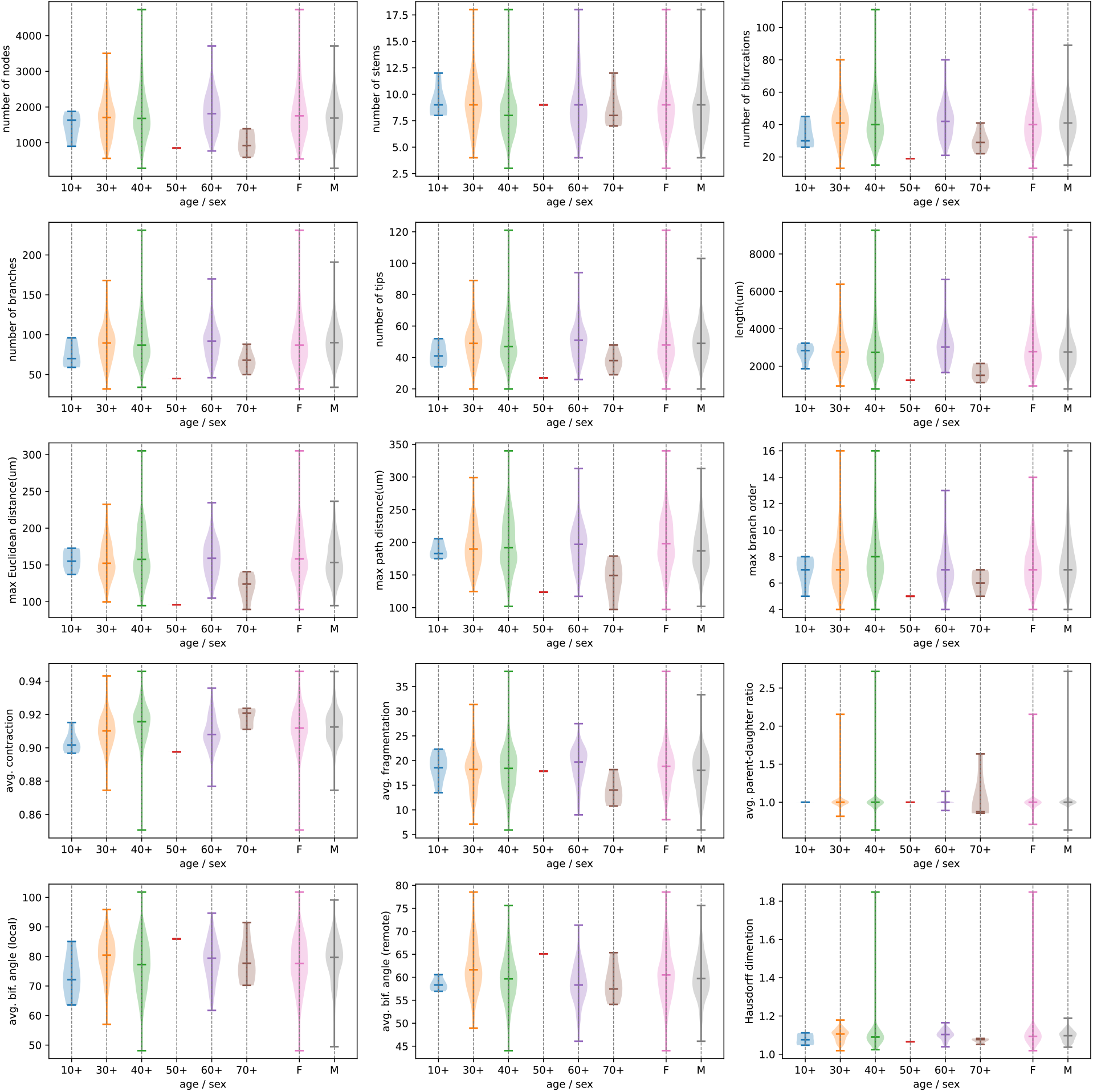
Exemplar global morphological features of each neuron reconstruction visualized as violin plots, grouped according to age or gender (F/M). The age was binned with a 10-year window to simplify the visualization. Each violin set shows the distribution of the values from a particular group, with the median value indicated by the bar in the middle.

**Supplementary Figure 7.**
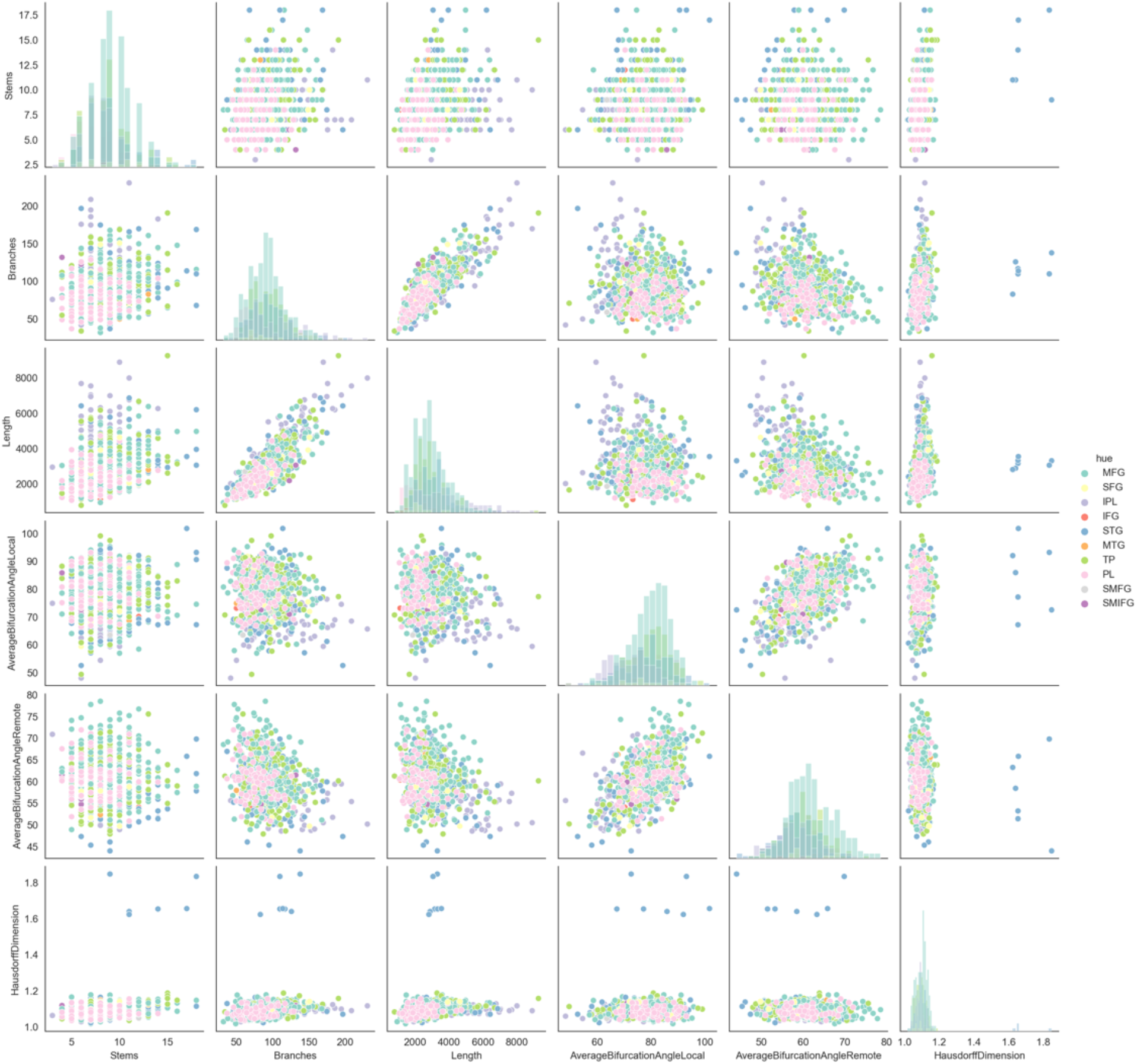
Pair-plot of 6 key global morphological features across 10 brain regions.

**Supplementary Figure 8.**
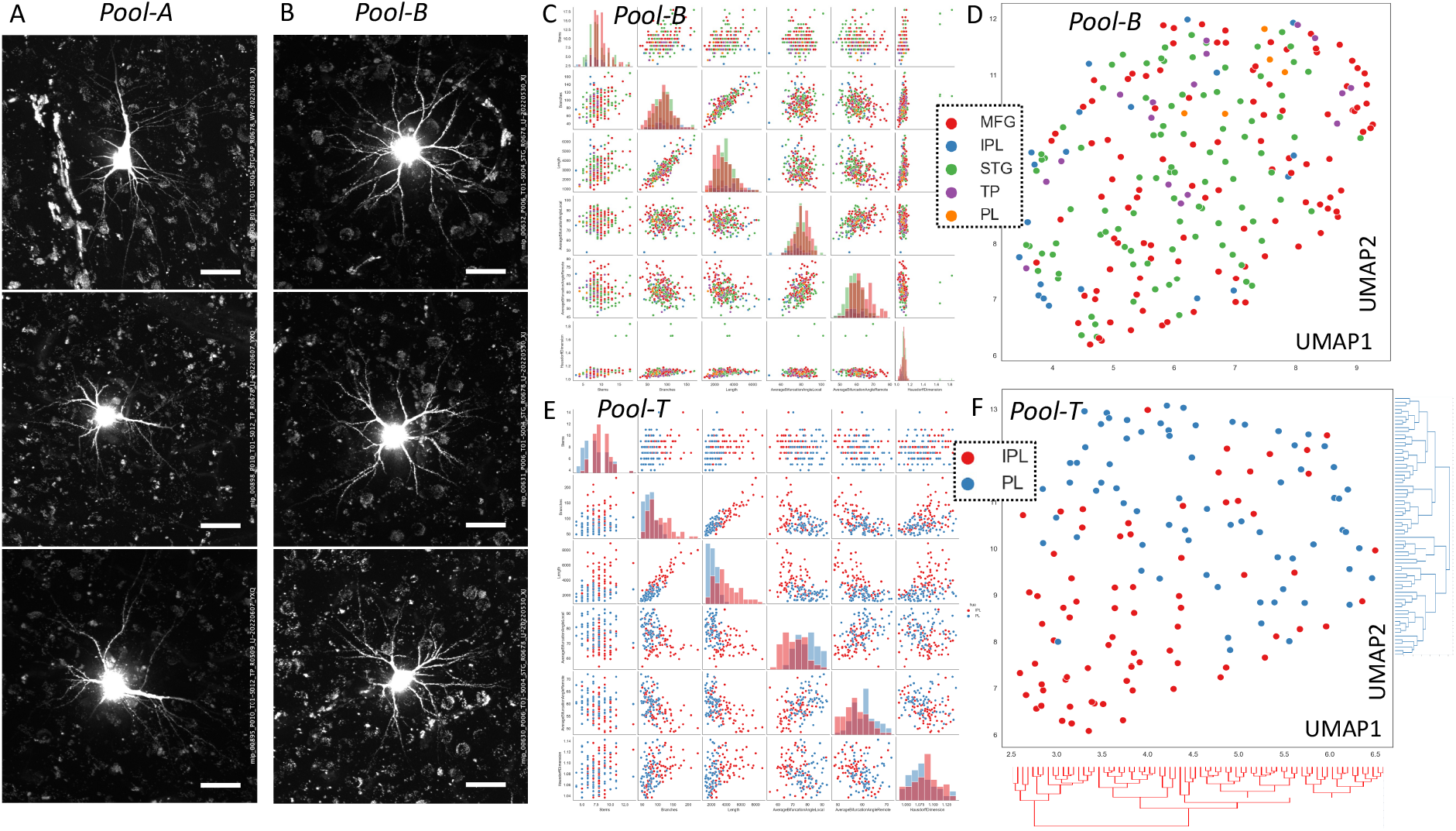
Orientation-classified analyses of interareal diversity of traced human neurons. A. Three examples of Pool-A neurons, where the apical dendrites align on the XY imaging plane. Scalebar is approximately 30μm. B. Three examples of Pool-B neurons, where the basal dendrites align on the XY imaging plane, perpendicular to the apical dendrites which align approximately along the Z-axial direction. Scalebar is approximately 30μm. C and D. Interareal pair-plot and UMAP analysis of 5 brain regions using 6 key morpho-features, in the same order and plotting convention in Figure 4A, but based on Pool-B neurons. E and F. Interareal pair-plot and UMAP analysis of IPL and PL neurons, using 6 key morpho-features, in the same order and plotting convention in Figure 4B, but based on Pool-T neurons. Dendrograms for IPL and PL neurons were also displayed in F to indicate the respective hierarchical clusters.

**Supplementary Figure 9.**
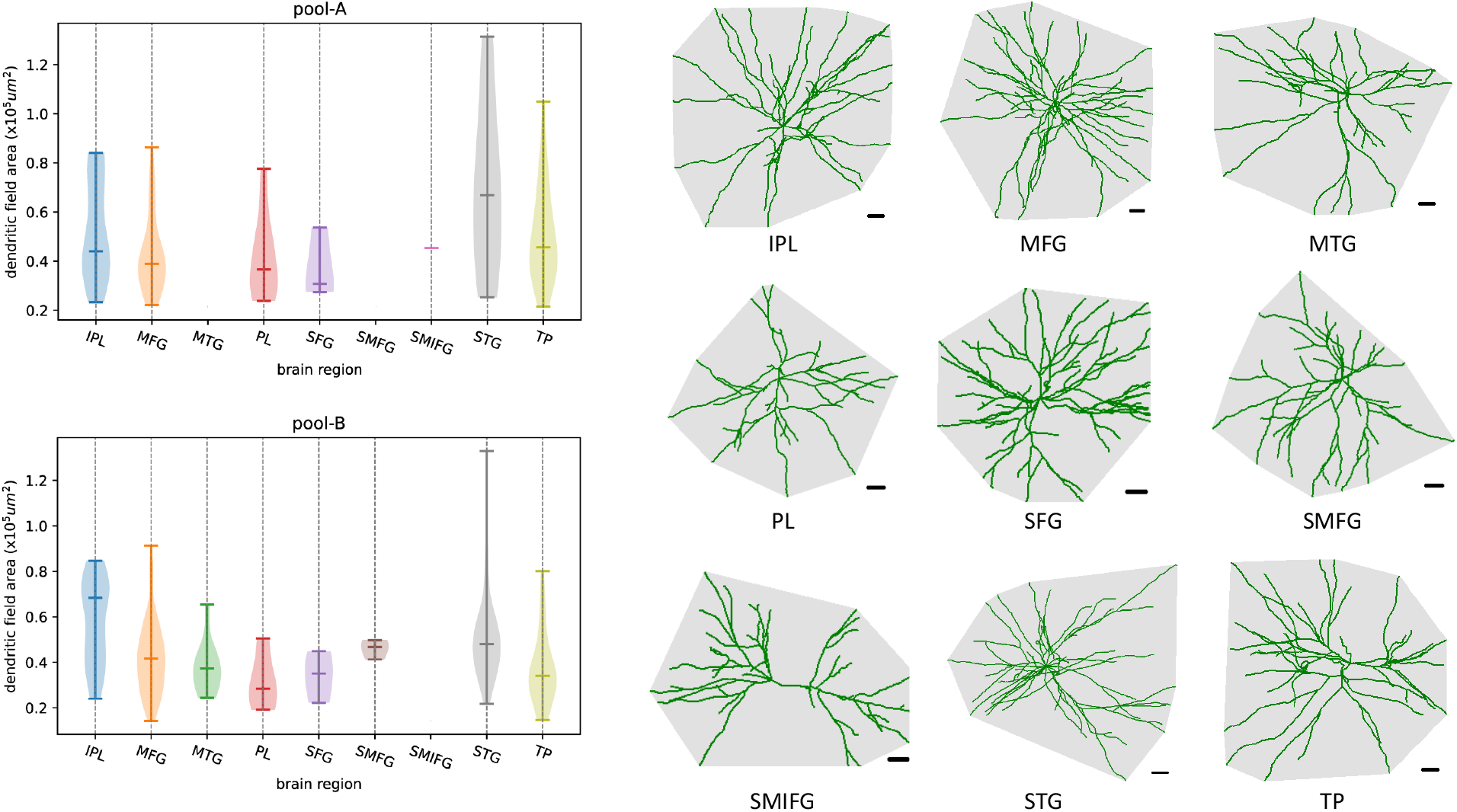
The results of the dendritic field area (DFA) for neuron reconstructions from pool-A and pool-B, shown as violin plots according to the brain regions. In particular, the DFA represents the area contained by the convex hull around the neurite region after being projected onto the XY-plane. Example neurite projections for each brain region from Pool-A (SMIFG) or Pool-B (other regions except SMIFG) are shown on the right of the figure along with the convex hull (scalebar=20um). Note that no violin plot was shown when the data in respective brain regions of Pool-A or Pool-B was absent.

**Supplementary Table 1.** Information of human brain tissue extraction and number of neurons imaged. Abbreviations: **SFG**: superior frontal gyrus; **MFG**: middle frontal gyrus; **IFG**: inferior frontal gyrus; **STG**: superior temporal gyrus; **MTG**: middle temporal gyrus; **PL**: parietal lobe ^**#**^; **IPL**: inferior parietal lobule; **AG**: angular gyrus; **TP**: temporal pole; **ACC**: anterior cingulate cortex; **ACG**: anterior central gyrus; *region involving SFG and MFG; **region involving SFG, MFG, and IFG. (L) and (R) indicate the left or right side of the respective brain. ^#^PL as used in this annotation was also suggested to be named as superior parietal lobe (Ding, et al, 2016), bordering with IPL. All the pathologies were diagnosed based on the fifth edition of the WHO Classification of Tumors of the Central Nervous System (CNS), published in 2021 (Louis DN, Perry A, Wesseling P, et al. The 2021 WHO Classification of Tumors of the Central Nervous System: a summary. Neuro-oncology 2021;23(8):1231-51. doi: 10.1093/neuonc/noab106 [published Online First: 2021/06/30]).

| Patient | Age | Sex | Diagnosis | Tumor/Disease Location | Neuron Specimen location | #Neurons (1746) |
| --- | --- | --- | --- | --- | --- | --- |
| H001 | 49 | M | Glioblastoma, IDH-wildtype, CNS WHO grade 4 | frontal/parietal lobes (L) | MFG near ACG | 48 |
| H002 | 36 | M | Glioblastoma, IDH-wildtype, CNS WHO grade 4 | frontal/temporal/insular/parietal lobes (L) | SFG (R) | 18 |
| H003 | 49 | F | Diffuse astrocytic glioma, H3-wildtype and IDH-wildtype, NOTCH1 mutant, Not Elsewhere Classified, CNS WHO grade 1 | thalamus (R) | IPL near AG | 225 |
| H004 | 71 | F | Glioblastoma, IDH-wildtype, CNS WHO grade 4 | temporal/insular lobes (R) | IFG | 4 |
| H005 | 36 | F | Diffuse hemispheric glioma, H3 G34-mutant, CNS WHO grade 4 | frontal lobe (R) | MFG | 335 |
| H006 | 63 | M | Glioblastoma, IDH-wildtype, CNS WHO grade 4 | temporal lobe (R) | STG | 171 |
| H007 | 35 | F | epidermoid cyst | middle cranial fossa (L) | MTG | 21 |
| H008 | 47 | F | Glioblastoma, IDH-wildtype, CNS WHO grade 4 | parietooccipital (L); corpus callosum | PL | 80 |
| H009 | 46 | M | Astrocytoma, IDH-mutant, CNS WHO grade 4 | frontal lobe (L); ACC (L); Corpus Callosum | SFG | 17 |
| H010 | 41 | M | Astrocytoma, IDH-mutant, CNS WHO grade 2 | temporal Lobe (L) | TP | 294 |
| H011 | 42 | F | Epilepsy | temporal Lobe (L); hippocampus (L) | STG | 31 |
| H012 | 28 | F | Subependymoma, CNS WHO grade 1 | lateral ventricles (R) | PL | 4 |
| H013 | 18 | M | Diffuse high-grade glioma, H3-wildtype and IDH-wildtype, RTK 1, CNS WHO grade 4 | frontal lobe (R); ACC (R); Corpus Callosum | S(M)FG* | 13 |
| H014 | 34 | M | Central neurocytoma, CNS WHO grade 2 | lateral ventricles (R) | MFG | 267 |
| H015 | 65 | F | Metastases to the brain and spinal cord parenchyma | frontal lobe (R) | S(M,I)FG** | 14 |
| H016 | 55 | M | Glioblastoma, IDH-wildtype, CNS WHO grade 4 | parietal lobe (L) | PL | 1 |
| H017 | 71 | M | Glioblastoma, IDH-wildtype, CNS WHO grade 4 | frontal/temporal/insular/parietal lobes (L); hippocampus (L) | IFG | 2 |
| H018 | 41 | F | Fibrous meningioma, CNS WHO grade 1 | lateral ventricles (L) | PL | 193 |
| H019 | 67 | F | Oligodendroglioma, IDH-mutant, and 1p/19q-codeleted, CNS WHO grade 3 | frontal pole (R); basal ganglia (R); corpus callosum | SFG | 8 |

